# Human prefoldin modulates co-transcriptional pre-mRNA splicing

**DOI:** 10.1101/2020.06.14.150466

**Authors:** Laura Payán-Bravo, Sara Fontalva, Xenia Peñate, Ildefonso Cases, José Antonio Guerrero-Martínez, Yerma Pareja-Sánchez, Yosu Odriozola-Gil, Esther Lara, Silvia Jimeno-González, Carles Suñé, Mari Cruz Muñoz-Centeno, José C. Reyes, Sebastián Chávez

## Abstract

Prefoldin is a heterohexameric complex conserved from archaea to humans that plays a cochaperone role during the co-translational folding of actin and tubulin monomers. Additional functions of prefoldin have been described, including a positive contribution to transcription elongation and chromatin dynamics in yeast. Here we show that prefoldin perturbations provoked transcriptional alterations across the human genome. Severe pre-mRNA splicing defects were also detected, particularly after serum stimulation. We found impairment of co-transcriptional splicing during transcription elongation, which explains why the induction of long genes with a high number of introns was affected the most. We detected genome-wide prefoldin binding to transcribed genes and found that it correlated with the negative impact of prefoldin depletion on gene expression. Lack of prefoldin caused global decrease in Ser2 and Ser5 phosphorylation of the RNA polymerase II carboxy-terminal domain. It also reduced the recruitment of the CTD kinase CDK9 to transcribed genes, and the association of splicing factors PRP19 and U2AF65 to chromatin, which is known to depend on CTD phosphorylation. Altogether the reported results indicate that human prefoldin is able to act locally on the genome to modulate gene expression by influencing phosphorylation of elongating RNA polymerase II, and thereby regulating co-transcriptional splicing.

## INTRODUCTION

Prefoldin is a protein cochaperone present in all living organisms except eubacteria (1). It is closely related to cytoskeleton assembly as it functions in the co-translational folding of actin and tubulin monomers (2, 3). While this first discovered function is cytoplasmic, prefoldin has also been found to play nuclear roles (4).

In eukaryotes, canonical prefoldin is composed of six different subunits, named PFDN1-6. Subunits PFDN2 and 6 are also components of the Uri/Prefoldin-like complex, involved in the cytoplasmic assembly of several protein complexes (5).

All the six yeast subunits are present in the nucleus and four of them (PFDN1, PFDN2, PFDN5 and PFDN6) are recruited to yeast chromatin in a transcription-dependent manner. Lack of these subunits cause transcription elongation defects, which is reflected in lower levels of long transcripts and genome-wide alterations in the profiles of active RNA polymerase II (RNA pol II) molecules (6). Yeast prefoldin mutants also show synthetic phenotypes when combined with mutations in transcription elongation and chromatin factors, and exhibit defects in co-transcriptional chromatin dynamics, indicating an important contribution of prefoldin to RNA pol II-dependent transcription and other phenomena linked to this process (6).

Human prefoldin subunits PFDN1, PFDN3 and PFDN5 have also been described to be involved in nuclear functions, including removal of HIV integrase (7), inhibition of c-Myc (8) and repression of some genes (9, 10). However, a possible general involvement of human prefoldin in transcriptional and co-transcriptional events has not been investigated. In this work we have explored this possibility and found a significant contribution of prefoldin to the expression of the human genome, particularly at the level of co-transcriptional mRNA splicing.

## MATERIALS AND METHODS

### Cell culture and transfection

Human HCT116 cells were maintained in McCoy’s 5A modified medium supplemented with 10 % foetal bovine serum, 60 mg/l penicillin, and 100 mg/l streptomycin. Additionally, 2 µg/µl G418 were added to resistant cell lines.

For transfection of 1 µg of plasmid or 15 µl siRNA 20 nM, 6 µl Turbofect transfection reagent (Thermo Fisher) and 8 µl Oligofectamine (Invitrogen) were used, respectively. Sequences of the siRNAs are as follows: siC (CGUACGCGGAAUACUUCGA); siPFDN2 (CAGCCUAGUGAUCGAUACA); siPFDN5 (AGAGAAGACAGCUGAGGAU).

### Generation of *PFDN5* null cell line by CRISPR-CAS9

The gRNA TTTCTTGGCTTTATATATCTTGTGGAAAGGACGAAACACCGGTACAGACCAAGTAT G was cloned by Gibson assembly following manufactureŕs instructions (Codex DNA) into the plasmid U6GRNA (Addgene 4148). 1 µg of the resulting plasmid and 0.5 µg of plasmid hCas9 (Addgene 41815) were co-transfected using 3 µl of Turbofect transfection reagent (Thermo Fisher). G418 resistant cells were cloned by dilution. *PFDN5* null mutant clones were selected by the absence of PFDN5 protein in western blot. A region of the *PFDN5* gene containing the gRNA target sequence was amplified by PCR, cloned into pGEMT (Promega) and sequenced.

### Plasmids and clone selection

*PFDN5* ORF (splice variant alpha) was cloned EcoRI/SalI into pCMV-Tag 2A (N-terminal Flag tag, Agilent) or XhoI/EcoRI into pEGFP-C1 (N-terminal GFP tag, Novopro). Expression of Flag-PFDN5 was assessed in the different clones by western blot, whereas in the case of PFDN5-GFP it was by flow cytometry.

### Antibodies

The following antibodies were used for chromatin immunoprecipitation (ChIP) or western blotting (WB): anti-PFDN5 (Santa Cruz, sc-271150, WB); anti-GAPDH (Santa Cruz, sc-47724, WB); anti-PFDN2 (Bethyl, A304-807A, WB); anti-Rpb1 CTD Ser2 phosphorylated (Abcam, ab5095, ChIP and WB); anti-Rpb1 CTD Ser5 phosphorylated (Abcam, ab5131, ChIP and WB); anti-Rpb3 (Abcam, ab202893, ChIP); anti-Rpb1 8WG16 (Thermo Fisher MA1-10882, ChIP); anti-FLAG (Sigma, F3165, ChIP); anti- U2AF65 (Santa Cruz, sc-53942, ChIP and WB); anti-PRPF19 (Lührmann Laboratory, ChIP and WB); anti-PRGL1 (Lührmann Laboratory, WB); anti-CDK9 (Santa Cruz, sc- 13130, ChIP and WB); anti-vinculin (Santa Cruz, sc-25336, WB).

### RNA-seq culture conditions

To analyse the possible functions of prefoldin in gene expression in human cells, we performed RNA-seq on siRNA transfected HCT116 cell line against *PFDN2* or *PFDN5* mRNA. The experiment began with the transfection of the cells, separately, with the different siRNAs. 24 hours later, the complete medium was replaced by medium without serum and the cells were incubated under these conditions for 48 hours. Finally, the serum-free medium was replaced by complete medium, and the samples were collected just before and 90 min after the addition of the serum.

### RNA extraction and quantitative RT-PCR assays

RNA was extracted with RNAeasy mini kit (Qiagen). 1 µg of this RNA was treated for 10 minutes at 65 °C with a unit of RQ1 RNAse-free DNase (Promega) in a total reaction volume of 10 µl. Next, it was retrotranscribed by Super-Script III (Invitrogen) using random hexanucleotides as primers. Relative quantification of specific cDNAs was achieved using SYBR premix (Takara) in a Lyght Cycler (Roche) and specific primers listed in Supplementary Table S1. The specific primer signal was normalized to the housekeeping gene GAPDH.

### RNA library preparation and sequencing

Total RNA was depleted of 18S rRNA and then libraries were prepared. 50 bp paired-end sequencing was performed at the Genomic Unit at CRG (Barcelona, Spain).

### RNA-seq analysis

Quality control was done with FSTQC v.0.11.5. Gene expresion was estimated using Kallisto v. 0.43 (11) against an index built on human genome GRCh38.p7 using -b 100 option. To calculate the exonic ratio, reads were mapped to the full human genome using HISAT2 v. 2.1.0 (12). Then exonic reads per gene were quantified using feature Counts v1.5.2 with options -p -B -O -t “exon” -g” “gene_id” using as annotation the file Homo_sapiens.GRCh38.87.gtf from emsembl v.87. Intronic read per gene were calculated with the same program and annotation file but with the following options: -p - B -O -t “gene” -g” “gene_id”. GO analysis was performed in R using GOFunction Package from Bioconductor. GO functions with an FDR<0.05 were selected as significant. To analyse how the length and number of introns of the genes affects the variation of the siPFDN5 / siControl log_2_ FC expression differences, genes with the highest expression level were excluded from this analysis (291 genes excluded out of 11700), since these are generally short (13, 14) and therefore could bias the results. Alternative Splicing Events were quantified using the SUPPA program(15).

### Co-transcripcional splicing assay

To investigate the rate of transcriptional elongation and pre-mRNA splicing, we grew cells on 60-nm plates to 70-80 % confluency and then treated them with 100 µM DRB (Sigma) for 3 hours. We washed the cells with PBS to remove the DRB and then incubated them in fresh medium for various time periods. After this incubation period, we lysed the cells and directly isolated total RNA. RT-qPCR with specific primers allowed us to measure, after the DRB wash, the time of appearance of newly synthesized pre-mRNA (forward primer on exon n, reverse primer on intron n + 1) and newly spliced pre-mRNA (forward primer on exon n, reverse primer on intron n + 2) (16).

### FAS Minigene assay

Cells were transfected with 0.5 µg of the reporter *FAS* minigene plasmid containing exons 5, 6 and 7 of the *FAS* gene (17). 24 hours later, RNA was extracted and retrotranscribed with MLV retrotranscriptase (Invitrogen) and a specific Fas primer (AAGCTTGCATCGAATCAGTAG). cDNA was amplified by 35 PCR cycles, DNA fragments were separated in an agarose gel and the bands quantified using Image Lab 6.0.

### Western blot

For whole cell extracts, cells pellets were lysed in Laemmli buffer 2X (125 mM Tris- HCl pH 6.8; 20 % glycerol; 4 % SDS; 0.2 M DTT and 1 % bromophenol blue). Samples were loaded at 100000 cells/lane and separated by electrophoresis under denaturing conditions using the Mini-Protean IITM system (BioRad) for 50 minutes at 180 V. A commercial protein mixture (Dual Color, BioRad) was used as a molecular weight standard. After finishing the electrophoresis, the proteins present in the polyacrylamide gels were transferred to nitrocellulose membranes at 90 V for 45 minutes. Next, membranes were blocked with 4% (w/v) skimmed milk in TTBS (15 mM Tris-HCl pH 7.5, 200 mM NaCl and 0.1 % Tween 20) for 1 hour at room temperature. Primary antibodies are listed in “antibodies”. Membranes were washed 3 times for 5 minutes in TTBS, incubated with HRP-conjugated secondary antibody in 4% (w/v) skimmed milk in TTBS and visualized using SuperSignal West Pico PLUS (Thermo Fisher Scientific).

### Chromatin immunoprecipitation

Human chromatin immunoprecipitation was carried out by fixation (10 minutes, 1 % formaldehyde, 37 °C) and quenching (5 minutes, 2 % glycine), cells were lysed in lysis buffer (50 mM Tris-HCl, 10 mM EDTA, 1% SDS, pH 8.1 and protease inhibitors). Samples were then sonicated (Bioruptor, Diagenode) with cycles of 30 seconds for 15 minutes and the extracts of 300 bp obtained were cleared by centrifugation. Sonicated chroamtin were incubated with 50 µl of magnetic dynabeads protein A o G (Life technology) coupled to the specific antibody. Immunoprecipitated material was washed with three different wash buffers (wash buffer 1: 0.1 % (w/v) SDS; 1 % (v/v) Triton- X100; 2 mM EDTA; 20 mM Tris-HCl pH 8; 150 mM NaCl. Wash buffer 2: 0.1 % (w/v) SDS; 1 % (v/v) Triton-X100; 2 mM EDTA; 20 mM Tris-HCl pH 8; 500 mM NaCl. Wash buffer 3: 0.25 M LiCl; 1 % (v/v) NP-40; 1 % (w/v) Sodium deoxicholate; 1 mM EDTA; 10 mM Tris-HCl pH 8) and TE (10 mM Tris-HCl pH 8; 1 mM EDTA). DNA was eluted using elution buffer (1 % (w/v) SDS; 0.1 M NaHCO_3_) and, treated with 5 µl proteinase K (20 mg/ml) for 1 hour at 37 °C. The resultant DNA was fenol-chloroform extracted, and precipitated. After resuspension in 200 µl of water, the DNA was quantified by qPCR. The levels of IP were normalized to input or negative control (material incubated without antibody). All primers used are listed in Supplementary Table S1.

To test whether prefoldin binds chromatin through nascent RNA, the fixation and sonication steps were reduced to 6 minutes. The sonicated chromatin was treated before immunoprecipitation with an RNase cocktail (300 U RNase T1 + 45 U RNase A per sample, Ambion) for 2 hours at 30 °C.

### ChIP-seq analysis

Libraries preparation and sequencing was performed at the Genomic Unit of CABIMER (Sevilla, Spain). Two biological replicates of the ChIP-seq data for each condition were primarily filtered using the FASTQ Toolkit program. Data were aligned using align function from Rsubread package, to map reads to the hg19 human reference genome using type = 1, TH1 = 2 and unique = TRUE parameters. The downstream analysis was performed on bamfiles with duplicates removed using the samtools rmdup command. Bigwig files were created on Flag-PFDN5 over Flag for each replicate using bamCompare from deeptools package (v3.1.3) with --scaleFactorsMethod readCount -- operation log2 parameters. Density plot was created using computeMatrix from deeptools package with -m 10000 -bs 50 -b 5000 -a 5000. EnrichedHeatmap (v1.12.0) package was used for drawing heatmaps for PFDN5 or RNA pol II (GSM1545657) on TSS +/- 2000 bp. For correlation plots, ChIP-seq signal was quantify on gene bodies or promoters (TSS +/- 500 bp) for PFDN5, RNA pol II (GSM1545657) or Ser2- phosphorylated RNA pol II (GSM4113565). Then all genes were sorted according its PFDN5 signal and grouped in 100 non-overlapping bins. The mean of each group was computed. Myc-dependent genes list was obtained from (18).

## RESULTS

### Transcriptomic analysis of PFDN5 and PFDN2 deficient human cells

In order to explore the potential role of prefoldin in human gene expression, we conducted experiments to analyse the impact of prefoldin perturbation on the cell transcriptome. We designed siRNAs for *in-vivo* depletion of subunits PFDN2 and PFDN5. PFDN5 is present in the canonical complex, whereas PFDN2 is a component of both the canonical and the Uri/prefoldin-like complex. We reasoned that those changes that are related to the canonical complex would be detected in the two experiments, whereas those related to the Uri/prefoldin-like would be found just in the PFDN2-depletion experiment.

Transfection of HCT116 human colon carcinoma cells with siRNAs that target *PFDN2* and *PFDN5* mRNAs (siPFDN2 and siPFDN5, respectively) had a 67 % and 55 % efficiency, respectively, in reducing the specific protein after 72 hours of treatment (Sup. Figure S1A).

In order to test the impact of prefoldin depletion on gene expression under different conditions of transcription regulation, we used serum stimulation after starvation, which has long been recognized as a fast and simple system to study global changes in transcription (19). We designed an experiment in which we transfected cells with the siRNAs for 24 hours, then serum-starved them for 48 hours, and finally stimulated by adding serum. Under these conditions, the efficiency of the siRNAs probed to be equally good (Sup. Figure S1B). Moreover, serum starvation seemed to lessen the impact on viability of the PFDN5 siRNA (compare Sup. Figure S1 B and D). We took samples just before and 90 minutes after stimulation in order to analyse transcriptomic changes by RNA-seq.

Global changes in RNA-seq signal were apparent in siPFDN5-treated cells both before and after serum stimulation (Figure 1A, left panels). The same was true for siPFDN2- treated cells, although in this case there were less differentially expressed genes (Figure 1A, right panels). In both cases, the number of significantly affected genes was higher after serum stimulation (Figure 1A, Sup. Figure S2A), suggesting that prefoldin may play a role in supporting regulated gene expression.

**Figure 1:**
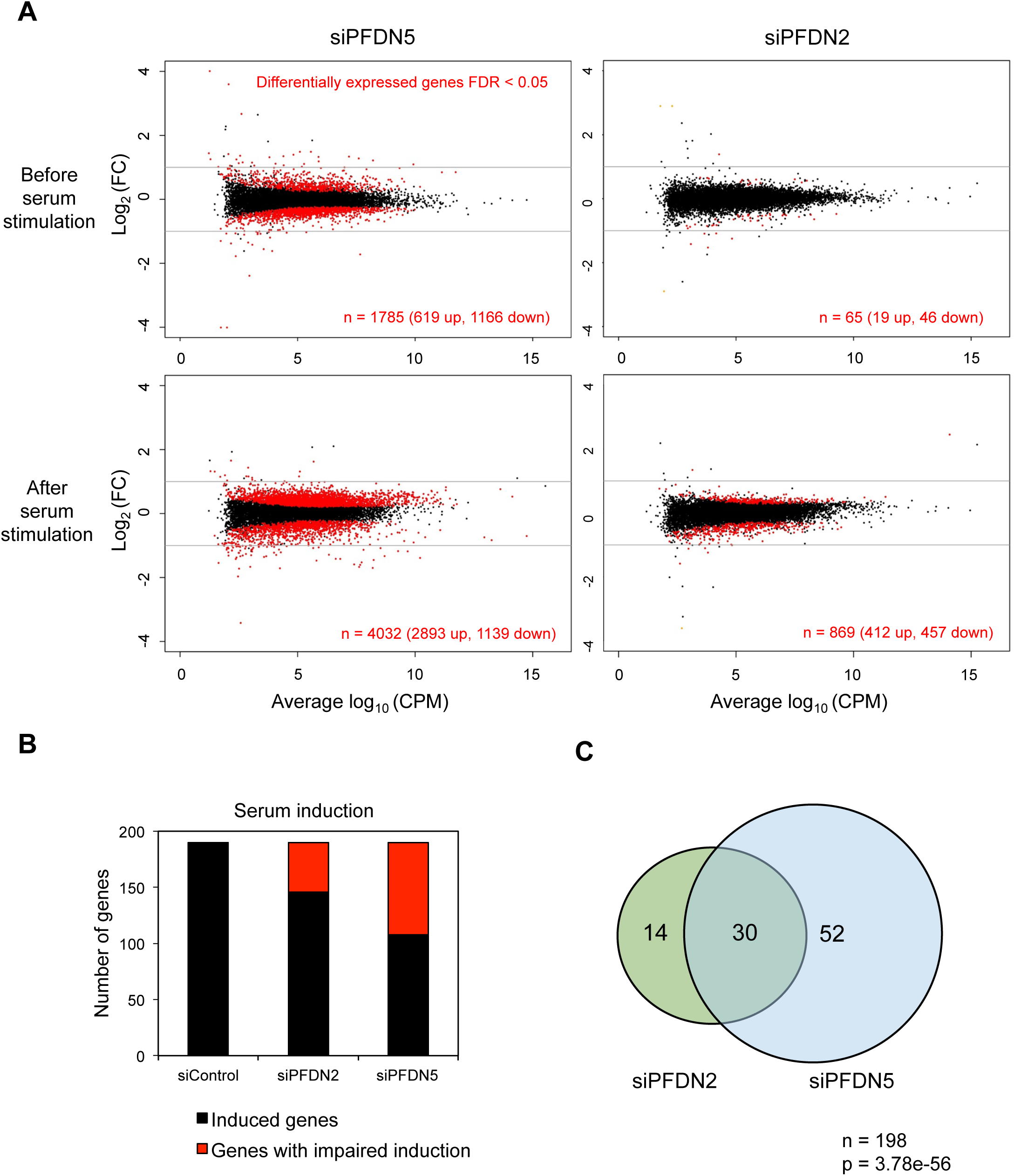
Two prefoldin subunits, PFDN5 and PFDN2, influence gene expression in human cells. HCT166 cells were treated with the different siRNAs for 24 hours, then serum-starved for 48 hours, then gene expression was induced by adding serum, and samples were taken just before and 90 minutes after stimulation. **A)** MA plots of RNA-seq experiments. The logarithm of the fold change (log_2_(FC siPFDN/siControl)) is shown against the logarithm (log_10_(CPM)) of the level of gene expression (defined for each gene as the mean of the counts per million of all the samples). The graphs on the left show the transcriptomic changes of the samples transfected with the siPFDN5 with respect to the control, and those on the right, those of the samples transfected with the siPFDN2. Unaffected genes are represented in black, while in red are all those genes showing |log_2_(FC)| > 0, FRD < 0.05. The upper panels represent the data of the samples taken before serum stimulation, while the lower panels represent those taken after serum stimulation. **B)** Representation of the number of genes induced by serum in control cells (log_2_(FC after/before serum treatment) > 1, FDR < 0.05), and those whose stimulation was significantly affected by siPFDN2 or siPFDN5 (FDR > 0.05). **C)** Overlap between those genes whose induction by serum was impaired by PFDN2 and PFDN5 depletion. The p-value after a hypergeometric test is also shown.

The higher levels of PFDN2 present in human cells, as compared to PFDN5 (20), may explain the lower impact of PFDN2 depletion. Nevertheless, gene ontology (GO) analysis of differentially expressed genes after serum stimulation showed significant enrichment of identical categories in both PFDN2- and PFDN5-dependent genes (Sup. Figure S2B). Before serum stimulation, despite a stronger effect of PFDN5 depletion, there was a significant overlap between the genes affected by PFDN5 and PFDN2 depletions (Sup. Figure S2C). Moreover, most genes affected in both knockdowns had the same direction of change (reduced expression in both or overexpression in both, Sup. Figure S2D), suggesting that most gene expression changes detected in PFDN5- and PFDN2-depleted cells are reflecting shared functions of the two subunits.

Next, we focused on the 198 genes that were more significantly induced (log_2_(FC) > 1, FDR < 0.05) by serum in control HCT116 cells (Figure 1B). Induction of 82 and 44 genes were significantly impaired in PFDN5- and PFDN2-depleted cells, respectively (Figure 1B). Comparison of these two subsets of genes resulted in a very significant overlap (Figure 1C).

Considered altogether, these results are compatible with a positive contribution of prefoldin to gene expression, particularly under conditions of gene regulation.

### Prefoldin deficiency causes stronger expression defects in long genes with a high number of introns

Yeast prefoldin has an impact on transcription elongation (6). If this were the case for human prefoldin, we would expect that longer genes would have more problems to be expressed properly in prefoldin knockdowns. To test this possibility, we divided the genes into quintiles according to their length and plotted the differential expression before and after serum stimulation. Before serum stimulation, in cells treated with siPFDN5 or siPFDN2, there was no sign of negative correlation between gene length and differential expression (Figure 2A, left panel and Sup. Figure S3A, left panel). If any, we detected a slight positive effect in very long genes in siPFDN5-treated cells that was not observed in siPFDN2-treated cells (Figure 2A and Sup. Figure S3A, left panel). In contrast, after stimulation we detected a consistent negative effect of gene length over gene expression in both siRNA treatments (Figure 2A and Sup. Figure S3A, right panel).

**Figure 2.**
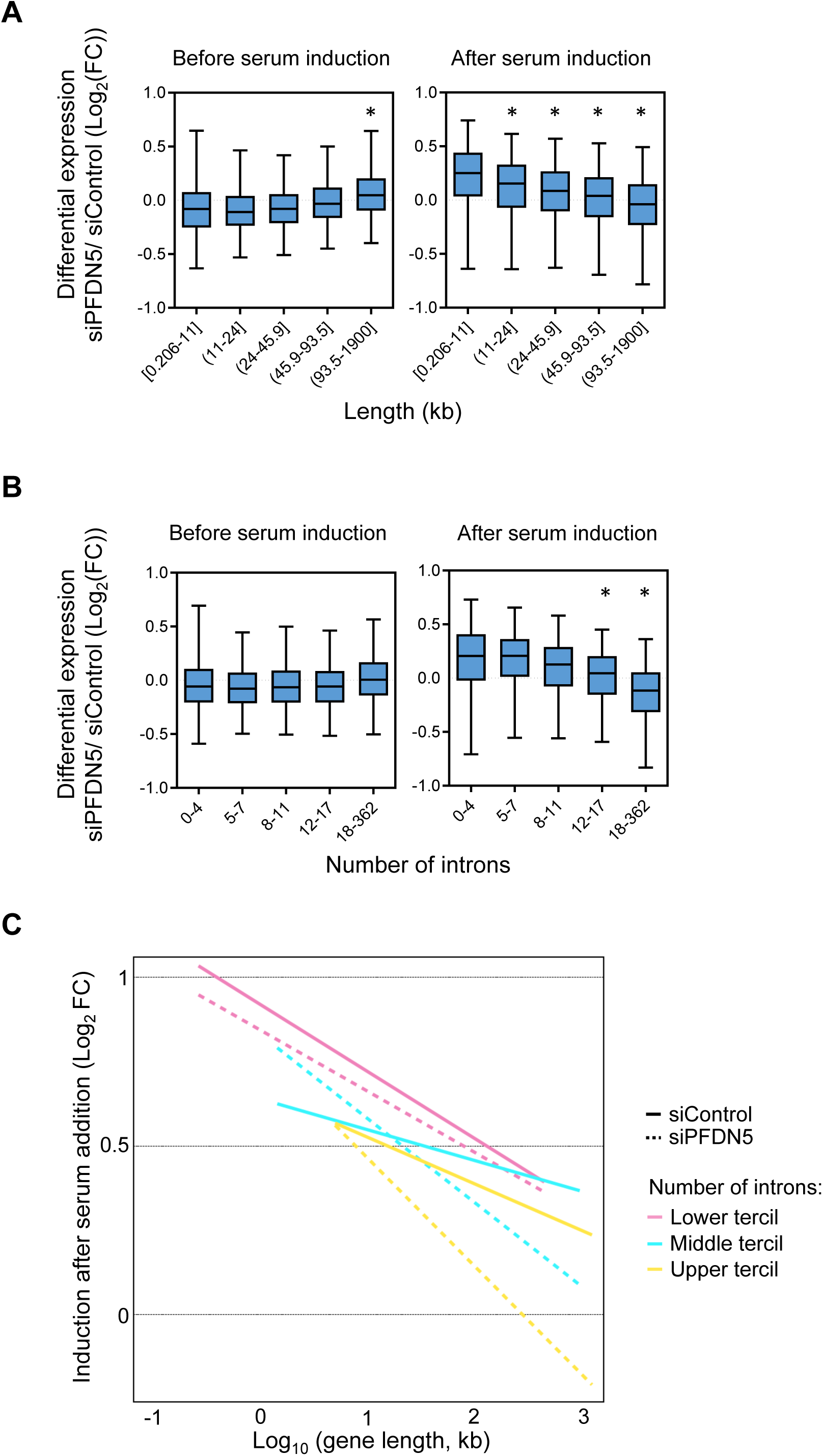
Prefoldin deficiency impairs the induction of long genes with a high number of introns. **A)** Expression differences between siPFDN5 and siControl cells (log_2_(FC)) with respect to the gene length, before and after serum stimulation. The genes were separated into quintiles. The * represents p < 0.005 in a Student’s t test comparing each quintile to quintile 1. **B)** The same analysis of A with respect to the number of introns that each gene contains, before and after serum stimulation. The genes were separated into quintiles. The * represents p < 0.005 in a Student’s t test comparing each quintile to quintile 1. **C)** Serum regulated genes (|log_2_(FC)| > 0 and FDR < 0.05, N = 1035) were divided into three groups according to the tercile of the number of introns. Serum-dependent fold change (Log_2_) is represented against the gene length (Log_10_), in cells transfected with the siControl (solid line) or siPFDN5 (dotted line).

In the human genome, gene length is proportional to the number of introns that a gene contains (21). Therefore, our next question was whether the number of introns correlated to the change in expression caused by prefoldin knockdowns. Genes were divided into quintiles depending on their number of introns. When differential expression of each group of genes was plotted, a negative relationship between expression and number of introns was apparent only after serum stimulation (Figure 2B, Sup. Figure S3B). These results are very similar to the previous ones, indicating an effect of the structural properties of genes (length, number of introns) in the expression under prefoldin deficiency conditions, pointing towards an effect of prefoldin in transcription elongation.

The effect of gene length could be a consequence of the effect of the number of introns, or viceversa. In a third possibility, both features could be independently contributing to the observed effect. To distinguish between these possibilities, we decided to plot the induction of the genes against their length. Since the two correlations were observed under serum stimulating conditions, we focused on those genes that were induced by serum in the control cells. These genes were further divided in three groups depending on their number of introns (Figure 2C). The negative effect of gene length in the capacity of a gene to undergo induction after serum addition was evident even in control cells (Figure 2C, Sup. Figure S3C). In these same control cells, the number of introns slightly enhanced the negative effect of length on the induction by serum. Strikingly, in the prefoldin knockdowns, there was a synergistic negative effect of length and intron number on gene induction, much stronger than in control cells (Figure 2C, Sup. Figure S3C). Taken altogether, these results point towards a possible influence of prefoldin in those events of the gene expression process, like pre-mRNA splicing, that take place or are set up during the elongation phase of transcription.

### Splicing efficiency after serum stimulation is reduced in prefoldin knockdown cells

To further investigate the possible role of prefoldin in splicing, we looked for defects in splicing in the RNA-seq data. Since polyA selection was not used for RNA-seq library preparation, we expected to find a certain, though low, level of unspliced mRNA precursors in the data set. In order to test this, we divided the reads from the sequencing reaction in two groups that we call “exonic reads” (those present in the mature mRNA) and “gene reads” (all the reads contained within genes, including those corresponding to unspliced precursors). For each gene, we divided the number of exonic reads by that of gene reads. Thus, we defined the exon ratio of each gene in each condition. The exon ratio is used here as a proxy for the maturation state of the products of a given gene in a given sample, with a ratio of 1 meaning that all the RNAs detected are matured mRNAs, and a ratio lower than 1 meaning that unspliced precursors were detected. As expected, most of the genes in our data presented a ratio close to 1, but other values of the ratio were clearly detectable (Sup. Figure S4). In control cells, the exon ratios increased after serum stimulation (Figure 3A and Sup. Figure S4). This increase was especially frequent in highly expressed genes (Figure 3A). However, in prefoldin siRNA treated cells, the exon ratio was higher than in control cells before serum stimulation (Sup. Figure S4), but decreased after that and, again, this decrease was particularly frequent in highly expressed genes (Figure 3A).

**Figure 3.**
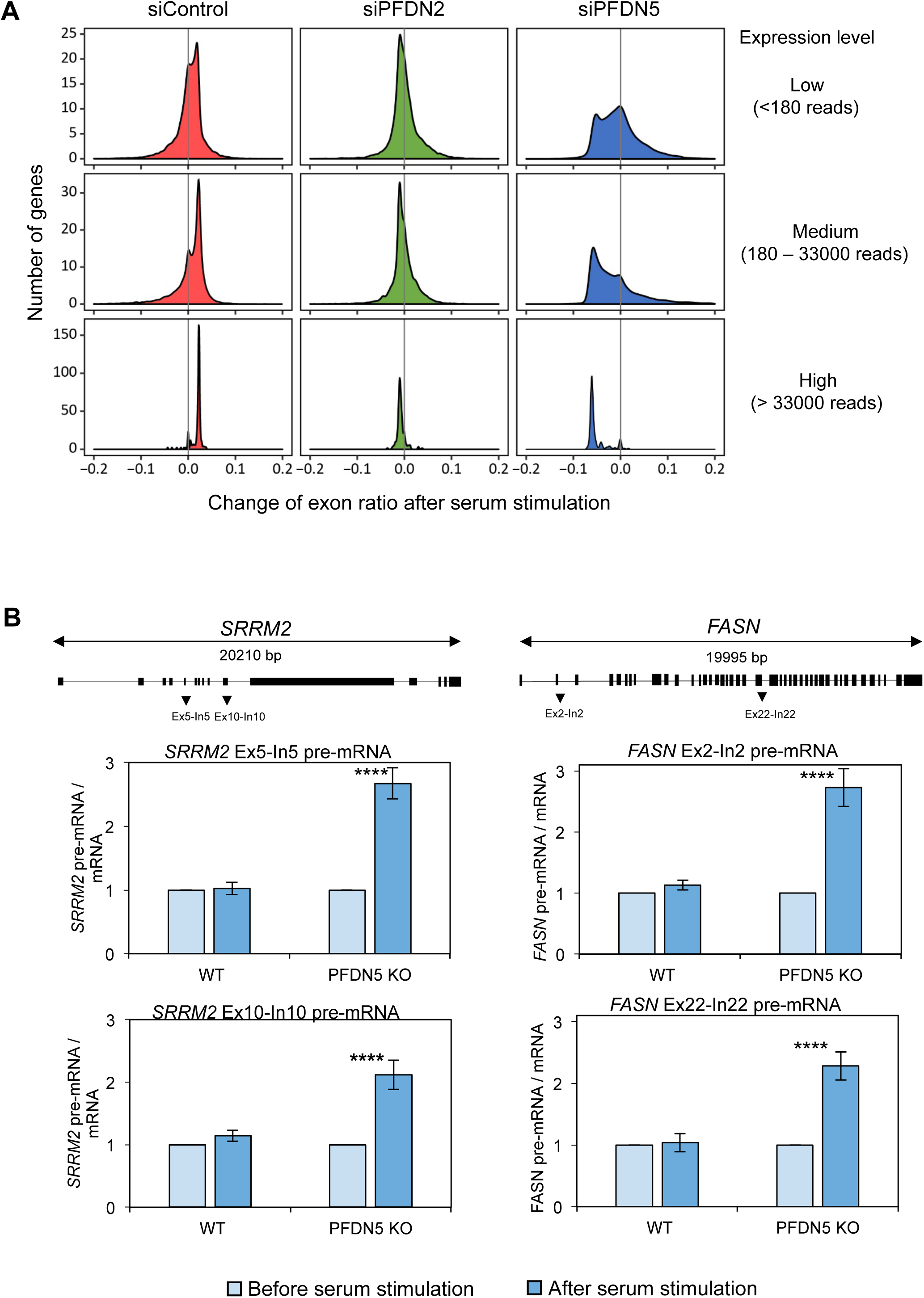
Prefoldin deficiency reduces splicing efficiency under serum stimulation conditions. **A)** The change of the exon ratio (exonic reads/ total reads) after serum stimulation is represented. The genes were divided into three groups according to their expression level (high, medium and low levels). The expression level of each gene was defined as the mean number of reads per million of all the samples, after normalizing by gene length. **B)** Control and *PFDN5* KO cells were treated as in Figure 1 and the level of pre-mRNA in two different regions of the *SRRM2* and *FASN* genes was determined by amplifying an intron-exon junction. The graphs represent the average and the standard deviation obtained from three different biological replicates. Pre-mRNA data was normalized first to mature mRNA level, and then to time zero. A total number of 3 biological replicates, with 3 technical replicates each, were considered for Student’s t test; ****p < 0.0001.

In order to exclude that these results were influenced by the stressful conditions of siRNA transfection, we validated them in *PFDN5* KO cells obtained by CRISPR-CAS9 (Sup. Figure S5A-C). Using our RNA-seq data, genes were ranked according to the difference in exon ratio in siPFDN5 treated cells before and after serum stimulation. Two model genes (*SRRM2* and *FASN*) were chosen among those with a higher difference. Levels of pre-mRNA for two introns of each gene were measured by RT-PCR in control and *PFDN5* KO cells. Higher levels of pre-mRNA in serum-stimulated cells were detected for all four introns in the mutant cells (Figure 3B).

Since pre-mRNA splicing impairment may affect alternative pre-mRNA processing, we analyzed different types of these events in the PFDN5 knockdown cells RNAseq data, by using the SUPPA computational tool (15). In about 20 % of the genes, we identified alternative processing events that were affected by the PFDN5 depletion, and found both suppressed (happening at least 10 % less frequently in PDN5-depleted than in control cells) and enhanced (happening at least 10 % more frequently in PDNF- depleted than in control cells). No class of event was particularly enhanced or suppressed by PFDN5 depletion (Sup. Figure S6A).

Exon skipping was one of the alternative splicing events analyzed by SUPPA. To further test the possible effect of prefoldin in alternative pre-mRNA splicing, we measured exon skipping in a transient transfection assay (minigene assay). *PFDN5* KO cells showed a mild increase in skipping of *FASN* exon 6 compare to control cells, and this increase was corrected by expressing an ectopic copy of *PFDN5* (Sup. Figure S6B).

These results suggest that the impairment of pre-mRNA splicing caused by prefoldin depletion may affect alternative processing, although in a very general manner, without introducing any specific bias in exon definition.

### Co-transcriptional splicing is delayed in prefoldin knockdown cells

Hitherto, we have shown evidence of human prefoldin being connected to pre-mRNA splicing and the expression of long genes. Since splicing can occur co-transcriptionally (22), the simplest explanation for this double connection of prefoldin would be that it is involved in co-transcriptional splicing. To test whether the prefodin-depleted cells were in fact defective in co-transcriptional splicing, we decided to do a DRB-mediated transcription inhibition and release experiment in siPFDN5-treated cells. DRB is an inhibitor of CDK9 that blocks transcription reversibly during early elongation (23). So, when DRB is washed out, polymerases start synchronously and the first round of transcription can be followed by RT-PCR. In order to study both transcription and co-transcriptional splicing, primer pairs were designed to amplify the cDNA derived from different kinds of unspliced precursors of the gene *CTNNBL1*, a long gene previously used as a model in this type of experiments (24, 25). We measured transcription elongation rate by the appearance of unspliced precursors with primer pairs spanning exon-intron junctions (Figure 4A). We could not detect any significant difference in the times required to detect exon 5-intron 5 or exon 6-intron 6 precursors. As expected, no signal of the intron 15-exon 16 precursor was detected at the same times (Figure 4A upper panels). Similarly, no delay was observed in the transcription of intron 19 of the *OPA1* gene (Sup. Figure S7A). When we repeated the same experiment in *PFDN5* KO cells, we did no find any change in the elongation rate of *CTNNBL1* (Figure 4A lower panels). These results ruled out a significant alteration of RNA pol II elongation rate by prefoldin depletion.

**Figure 4.**
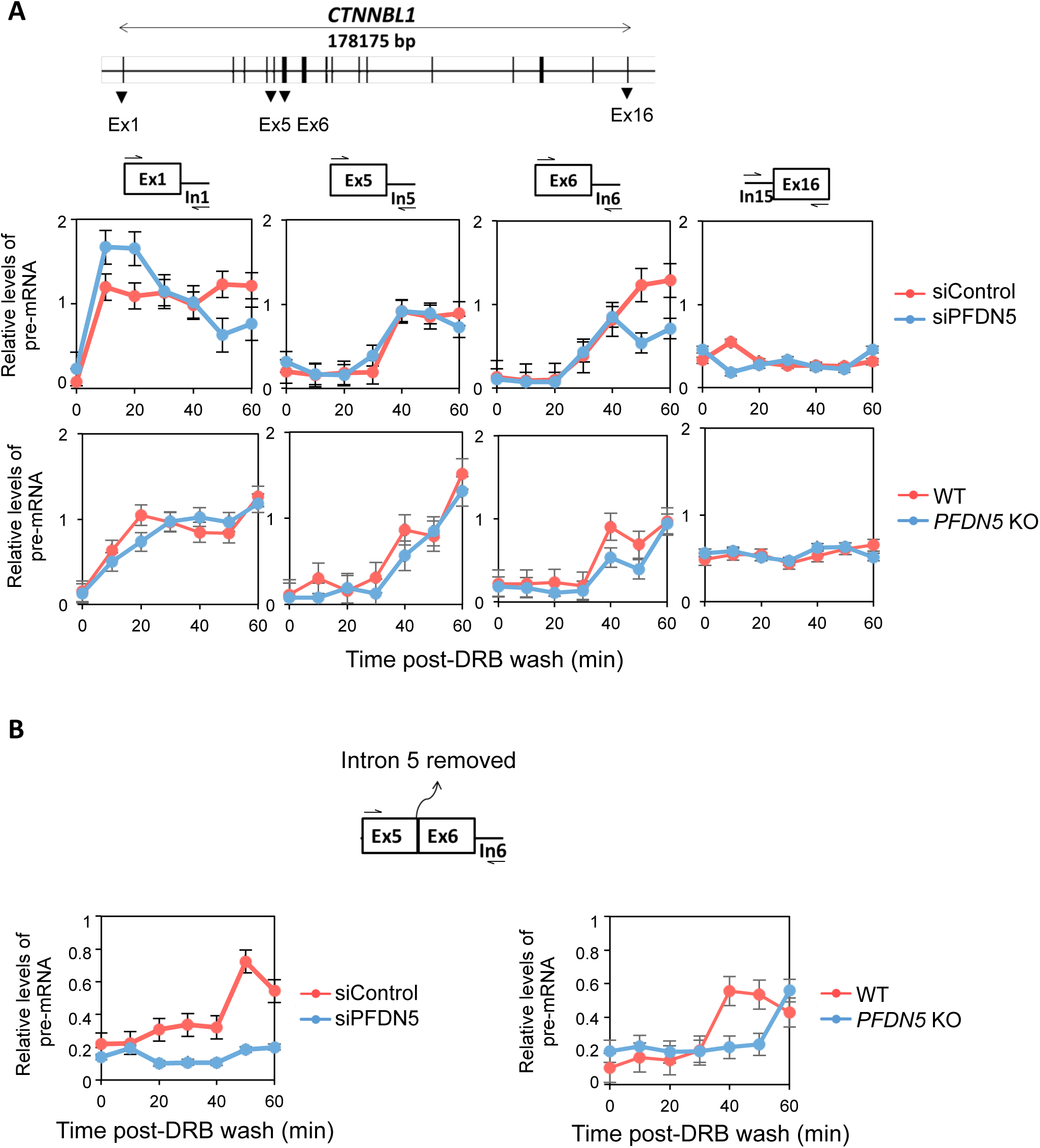
PFDN5 depletion impairs co-transcriptional splicing in the long *CTNNBL1* gene. **A)** Pre-mRNA levels at different loci throughout the *CTNNBL1* gene in cells transfected with control siRNA (red line) or PFDN5 siRNA (blue line) (upper panels), and in WT (red line) or *PFDN5* KO cells (blue line) (lower panels). Cells were treated with 100 µM DRB for 3 hours. Samples were taken every 10 minutes after washing DRB out. Pre-mRNA data was normalized first to the mature mRNA, and next to the DRB untreated sample. A scheme of the *CTNNBL1* gene is also shown, and the different amplicons used are depicted above each graph. **B)** Event of co-transcriptional splicing (measured as the presence of pre-mRNA without the indicated intron) in the *CTNNBL1* gene in cells treated as in A). The amplicon used is depicted above the graph. The average and standard error of at least three biological replicates are represented.

Using the same samples, we measured co-transcriptional splicing by the appearance of precursors containing intron 6 of *CTNNBL1*, but lacking intron 5, already spliced (Figure 4B). While in control cells this kind of precursors were detectable 50 minutes after DRB wash, in siPFDN5-treated cells we could not detect it before the end of the experiment (Figure 4B left panel). In *PFDN5* KO cells, we detected the same precursor with a 20 minutes delay compared to WT cells (Figure 4B right panel). In the case of the OPA1 genes, we also detected a delay in the detection of a pre-mRNA precursor containing intron 19 but lacking intron 18, when we compared PFDN5-depleted and non-depleted cells (Sup. Figure S7B). These results indicate that prefoldin is required for timely coupling between transcription elongation and pre-mRNA splicing.

### Prefoldin acts locally in the body of transcribed genes

We then explored whether the functional involvement of prefoldin in co-transcriptional splicing correlated with its physical presence in the transcribed gene. Prefoldin localizes to the nucleus of different organisms (6, 26–29) and, in yeast, it has been found associated to the body of transcribed genes (6). To test this in human cells by ChIP, a Flag-PFDN5 stable cell line was constructed (Sup. Figure S8A). When we examined the presence of Flag-PFDN5 in chromatin by ChIP-seq, we found it accumulated mainly around the transcription start site (TSS), following a distribution similar to RNA pol II (Figure 5A-B). Moreover, the signal was stronger in expressed genes compared to non-expressed ones (Figure 5A). PFDN5 signal was also detected in gene bodies, where it highly correlated with total RNA pol II and with its phosphorylated Ser2-CTD form (Ser2P) (Pearson correlation 0.84 and 0.88 respectively, Figure 5C). We also found significant binding of PFDN5 in the gene bodies of two long model genes by ChIP and Q-PCR (*CTNNBL1* and *CD44*, Figure 5D).

**Figure 5.**
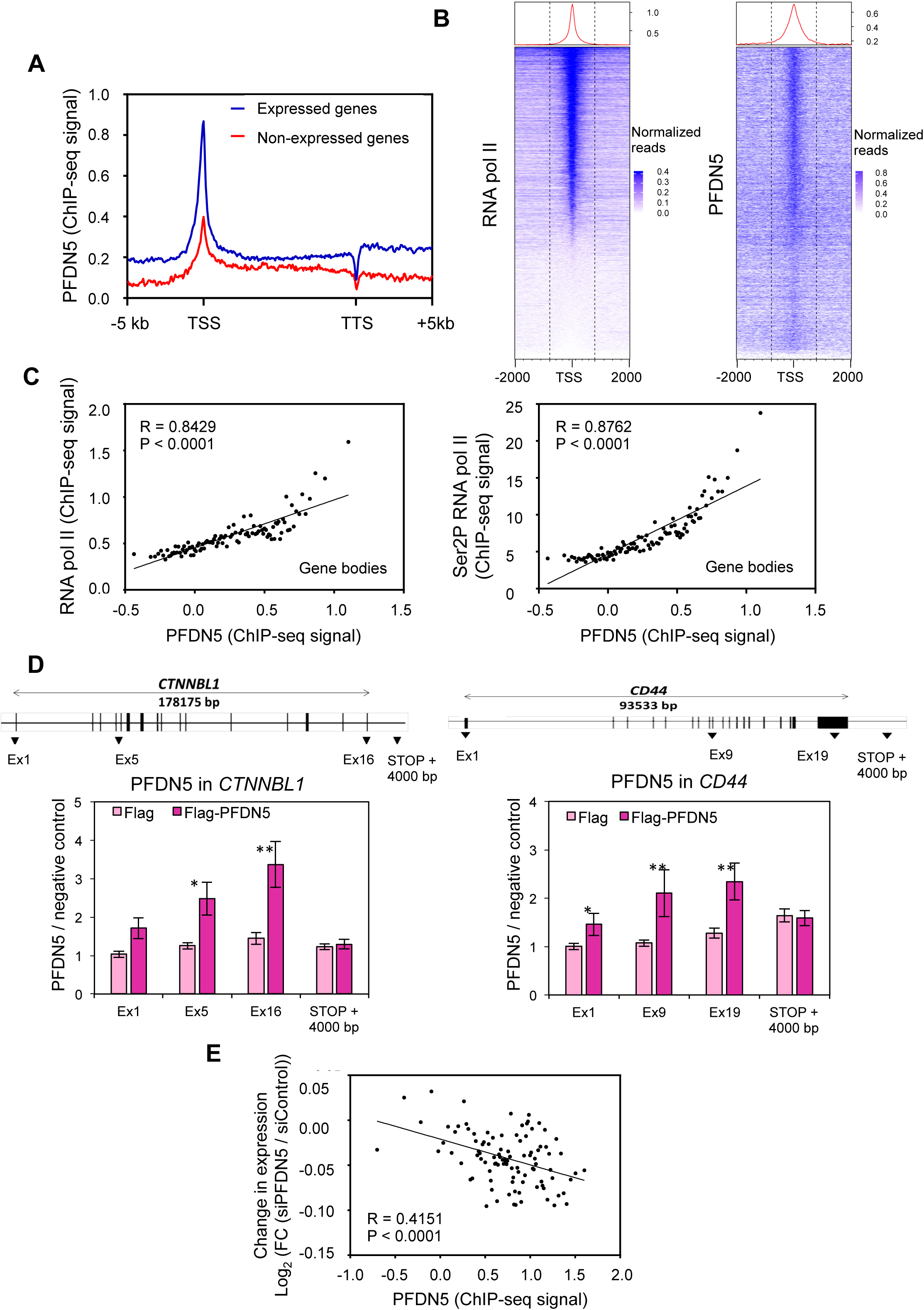
PFDN5 is present in the chromatin of active genes. **A)** Metagene analysis of the ChIP-seq signal of Flag-PFDN5. Protein coding genes were separated between expressed (>0.1 reads per kilobase per million, RPKM) and not expressed genes (<0.1 RPKM). Genes were scaled to the same length from the transcription start site (TSS) to the transcription termination site (TTS). Upstream and downstream sequences up to 5 Kb are represented unscaled. **B)** Heatmaps of the ChIP-seq signals of RNA pol II and PFDN5. Genes were ordered according to their RNA pol II signal. Centred in the TSS, 2 Kb are represented upstream and downstream. **C)** The gene body signal of total RNA pol II (left panel) or Ser2P RNA pol II (right panel) was represented with respect to the PFDN5 signal in the same region. Expressed genes (>0.1 RPKM) were ordered according to the PFDN5 signal and divided into 100 bins. The mean signal of each group was then calculated and represented. The R and p values from Pearsońs correlation are shown. **D)** The levels of Flag-PFDN5 were measured by ChIP-qPCR in different regions of the *CTNNBL1* (left panel) and *CD44* (right panel) genes using anti-Flag antibody in control (light pink) and Flag-PFDN5 cells (dark pink). An intergenic region in chromosome 5 was used as the non-transcribed negative control. A total number of 3 biological replicates, with 3 technical replicates each, were considered for Student’s t test. *p < 0.05, ** < 0.005. **E)** The PFDN5 signal was represented with respect to the fold change of expression in the siPFDN5 RNA-seq data. Expressed genes (>0.1 RPKM) were ordered according to the FC and divided into 100 bins. The mean signal of each group was calculated and represented against the mean PFDN5 signal around the promoter (TSS +/- 500 bp). The R and p values from Pearsońs correlation are shown.

Cell extracts pretreated with RNase before performing the immunoprecipitation step still showed significant ChIP signals, indicating that binding of PFDN5 to chromatin is not mediated by nascent pre-mRNA (Sup. Figure S8B).

The results described above confirm the presence of PFDN5 in the transcription sites across the human genome. However, this physical coincidence between prefoldin and transcribing RNA pol II does not demonstrate an active role of prefoldin in transcription, mediated by its location on transcribed genes. To test this possibility we compared PFDN5 ChIP signals to the functional effect of PFDN5 depletion on gene expression under serum stimulation conditions (previously described in Figure 1A and Sup. Figure S2A). We found a negative correlation between the PFDN5 ChIP-seq signal and the effect of PFDN5 depletion in gene expression (Figure 5D). In other words, the more PFDN5 in a gene, the more dependent its expression on PFDN5. We found this result very meaningful, considering that the correlation between the PFDN5 ChIP-seq and gene expression (RNA-seq) signals in non-depleted cells was weak (Sup. Figure S9). Taken altogether, these results indicate that PFDN5 is physically present in transcribed genes and that this presence has a positive local impact on gene expression.

### Prefoldin contributes to the recruitment of splicing factors to elongating RNA pol II

One possible mechanism on how prefoldin might influence co-transcriptional splicing is by favouring the action of splicing factors. In order to search for such a potential functional interaction between prefoldin and pre-mRNA splicing factors we made use of TheCellMap (https://thecellmap.org/). This database contains genetic interaction data for thousands of yeast mutants. Considering that yeast prefoldin has been also previously connected to transcription elongation (6), we reasoned that yeast genetic interactions might be informative for those functions of prefoldin that has been conserved during the evolution of the eukaryotic kingdom. When we systematically looked for correlations between the six prefoldin subunits and splicing factors, we obtained the results shown in Supplementary figure S10A. Most of the correlations found pointed to a functional interaction between prefoldin and the PRP19 complex (PRP19C). Indeed, PRP19C has been reported to be recruited to transcribed chromatin through interaction with the splicing factor U2AF65 in a CTD phosphorylation-dependent manner (30). Therefore, we decided to test the recruitment of PRP19 and U2AF65 to transcribed genes in *PFDN5* KO and siPFDN2 cells. Under conditions of prefoldin perturbation, both PRP19 and U2AF65 levels were reduced in the two genes tested as compared to control cells (Figure 6A-B, Sup. Figure S11A-B).

**Figure 6.**
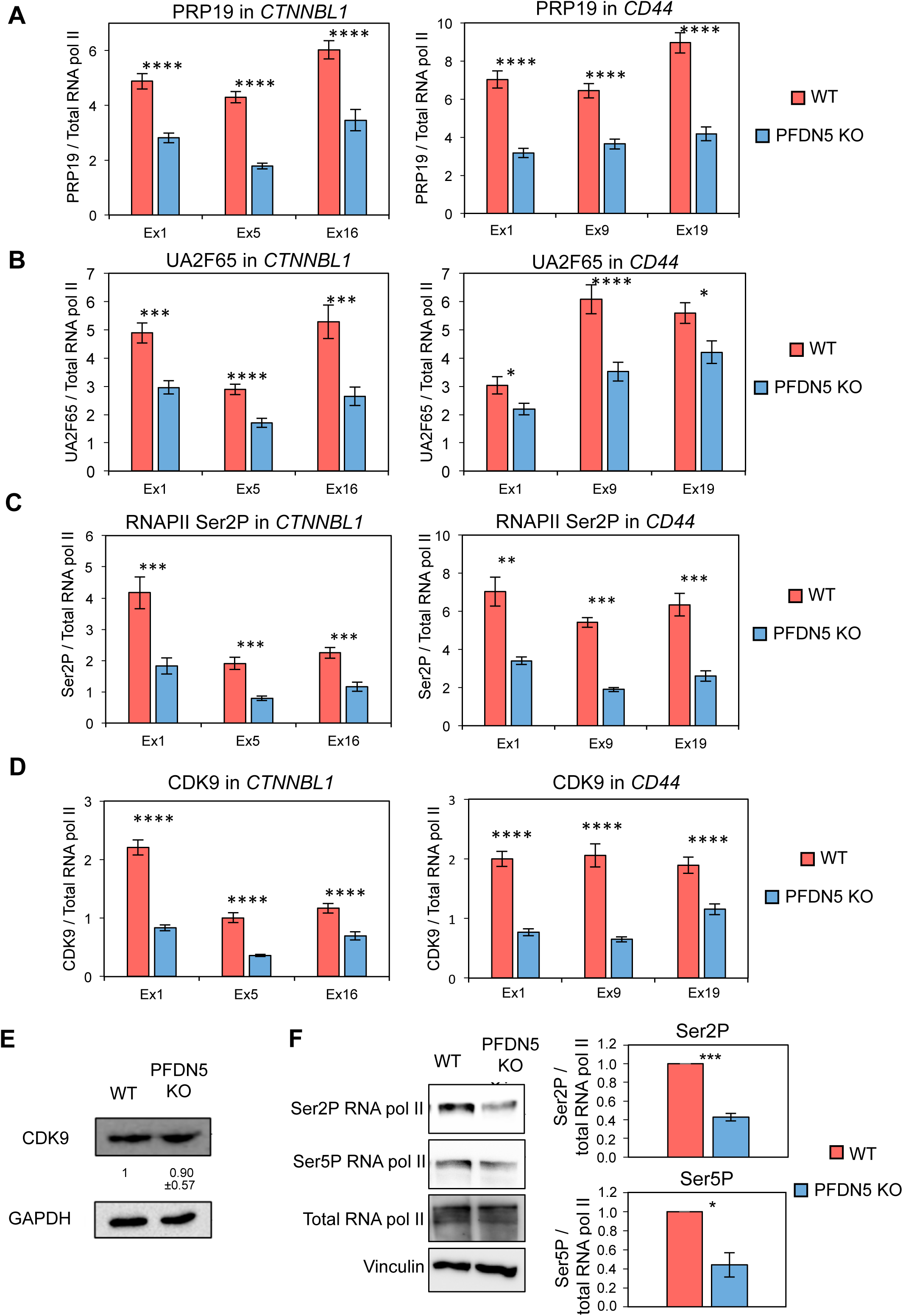
Lack of PFDN5 decreases PRP19, U2AF65 and CDK9 recruitment to transcribed chromatin and Ser2 phosphorylation of elongating RNA pol II. **A)** The levels of PRP19 were measured by ChIP in different regions of the *CTNNBL1* (left panel) and the *CD44* (right panel) genes, using anti-PRP19 antibody in control (red bars) and *PFDN5* KO cells (blue bars). Values were normalized to total RNA pol II levels. **B)** The levels of U2AF65 were measured by ChIP in different regions of the *CTNNBL1* (left panel) and the *CD44* (right panel) genes, using anti-U2AF65 antibody in control (red bars) and *PFDN5* KO cells (blue bars). Values were normalized to total RNA pol II levels. **C)** The levels of Ser2-phosphorylated RNA pol II were measured by ChIP in different regions of the *CTNNBL1* (left panel) and the *CD44* (right panel) genes using an anti-Ser2P-CTD specific antibody, in control (red bars) and *PFDN5* KO cells (blue bars). RNA pol II CTD-Ser2P values were normalized to total RNA pol II levels. **D)** The levels of CDK9 were measured by ChIP in different regions of the *CTNNBL1* (left panel) and the *CD44* (right panel) genes using an anti-CDK9 specific antibody, in control (red bars) and *PFDN5* KO cells (blue bars). Values were normalized to total RNA pol II levels. **E)** Global CDK9 protein levels analysed by western blotting in control and *PFDN5* KO cells. GAPDH was used as a loading control. Averaged values and standard deviation of the CDK9/GAPDH ratio from three experiments are shown. **F)** The global levels of Ser2P and Ser5P forms of RNA pol II were measured by western blot in control and *PFDN5* KO cells. Anti-vinculin antibody was used a loading control. A representative experiment is shown on the left. Average values and standard error of at least three biological replicates are shown in the graphs. * p-value < 0.05; ** < 0.005; *** < 0.0005; **** < 0.0001. A total number of 12 technical replicates were considered for Student’s t test.

No reduction was found for total RNA pol II levels (Sup. Figure S12A-B). Moreover, total protein levels of U2AF65 and several components of PRP19C remained constant in the absence of PFDN5 (Sup. Figure S13A-C).

In yeast cells, lack of PFDN1 causes a defect in Ser2 phosphorylation of RNA pol II CTD (6). CTD phosphorylation is known to favour the recruitment of some splicing factors (31) and, among them, PRP19 and U2AF65 (30). Thus, if the CTD phosphorylation defect were conserved in humans, we reasoned that reduced phosphorylation levels could result in impairment of U2AF65 recruitment, which in turn would originate co-transcriptional splicing problems and the kind of splicing defects that we observed in our RNA-seq data. Indeed, when we measured the level of Ser2P by chromatin immunoprecipitation in human *PFDN5* KO cells, we observed a significant reduction of the signal (Figure 6C).

CDK9 is the mayor kinase that controls Ser2P (32). Interestingly, we could observe a reduction in the levels of CDK9 bound to chromatin of the two genes tested in *PFDN5* KO cells (Figure 6D). Lack of prefoldin did not produce significant changes in the total cellular levels of CDK9 (Figure 6E).

Reduction of RNA pol II CTD phosphorylation in *PFDN5* KO cells was a general phenomenon across the genome, since western blot experiments showed lower global levels of Ser2P and also, although less significantly, of Ser5 phosphorylation (Ser5P) (Figure 6F).

Taken altogether, these results indicate that prefoldin contributes to co-transcriptional splicing by preserving RNA pol II phosphorylation and favoring the recruitment of PRP19C to transcribed genes in human cells.

## DISCUSSION

Prefoldin is well known for its cytoplasmic role in the co-translational folding of tubulin and actin monomers (33, 34). Prefoldin has also been shown to play additional functions in the cell nucleus (4, 35), including a positive role in transcription elongation and co-transcriptional chromatin dynamics across the yeast genome (6). The results described in this work extend this view as they show a general contribution of prefoldin to human gene expression by modulating co-transcriptional splicing. The detailed analysis of the transcriptome of prefoldin-depleted human cells that we have performed uncovered significant alterations of pre-mRNA splicing, lower levels of expression in genes with a high number of introns, and impaired induction of those serum-activated genes that contain a higher number of introns. Moreover, gene length and number of introns synergistically impaired gene induction by serum in prefoldin-depleted cells.

These data points towards a positive influence of prefoldin in co-transcriptional pre-mRNA splicing. Our experiments analysing DRB-synchronized populations of RNA pol II molecules confirmed that prefoldin-depleted cells were defective in co-transcriptional splicing.

We found that prefoldin binds transcriptionally active chromatin across the human genome and that prefoldin perturbation provokes gene expression defects and impaired pre-mRNA splicing. The simplest hypothesis to explain this set of phenomena involves that prefoldin would act locally during transcription elongation and that this action would favour gene expression by enhancing co-transcriptional pre-mRNA splicing. The negative correlation that we found between PFDN5 ChIP signal across the genome and the impact of PFDN5 depletion on gene expression (Figure 5E) lends solid support to this local action of prefoldin during transcription.

Our results suggest that prefoldin influences co-transcriptional pre-mRNA splicing by favouring the recruitment of the splicing machinery during transcription elongation. Pre-mRNA splicing is a sequential process that consists of spliceosome assembly, pre-catalytic activation and catalysis (36). It is known that spliceosome assembly is coupled to transcription elongation (37, 38). We found that genetic interactions point to a closer correlation of prefoldin subunits with the PRP19 complex, a splicing factor involved in the pre-catalytic activation step (Supplementary Figure S10B) (39), than with the snRNP components of the spliceosome (Supplementary Figure S10A). Similar level of correlation was also found between several prefoldin subunits and the splicing factor PRP2 (Supplementary Figure S10A), which also acts during remodelling of the spliceosomal Bact complex to the catalytically activated B* complex, just before step one of splicing (40). The action of PRP19C is mediated by U2AF65, which helps recruit this complex to the phosphorylated RNA pol II CTD domain (30). This mechanism explains coupling of transcription elongation and pre-mRNA splicing activation (30). We found that lack of prefoldin provokes a global decrease in RNA pol II CTD-Ser2P and Ser5P. The drop in Ser2P caused by prefoldin perturbation is consistent with the significant reduction in the levels of CDK9 found by ChIP. We also found a parallel decrease in the levels of PRP19 and U2AF65 present in the body of transcribed genes. We, therefore, propose a scenario where the positive contribution of prefoldin to CTD phosphorylation is necessary for the recruitment of PRP19C/U2AF65, which in turn allows efficient co-transcriptional splicing (Supplementary Figure S14) (30). We cannot exclude that prefoldin, by favouring the recruitment of U2AF65 to elongating RNA pol II, is also contributing to transcription elongation itself. In fact, Mud2, the yeast U2AF65 homolog, favours both transcription elongation and pre-mRNA splicing (41). However, we did not find significant reduction in the elongation rate by RNA pol II in the two genes tested.

Another, non-mutually exclusive possibility is that canonical prefoldin helps the assembly of the spliceosome. The co-chaperone function of prefoldin could be necessary for this. Other large complexes are assisted by the Uri/prefoldin-like complex for their assembly (5, 42, 43). Moreover, the prefoldin-like component RUVBL1/RUVBL2 regulates assembly of the U5 small nuclear ribonucleoprotein (42), and plant canonical prefoldin is involved in maintaining the levels of the LSM2-8 complex through Hsp90 (44). However, this potential role of human prefoldin cannot explain all the results presented here, since an assembly failure would affect spliceosome activity in general, not only co-transcriptional splicing.

Prefoldin is a stable heterohexameric complex (20). However, there is also evidence that some of the individual subunits have a function by themselves (45). Human PFDN5, for instance, is able to inhibit the transactivation domain of c-Myc by recruiting histone deacetylases (9) and promoting c-myc degradation (46). This interaction of c-Myc and PFDN5 is subunit-specific and does not involve other prefoldin subunits (9). However, most binding of prefoldin to chromatin does not seem to depend on Myc. Although slightly lower in average than in Myc target genes, PFDN5 ChIP signals were also clearly detectable in non-target genes, in spite of the lower average transcription levels of the latter (Sup. Figure S15A-B). Moreover, we detected clear parallelisms between PFDN2 and PFDN5 in the analyses that we performed. We conclude that the effects on co-transcriptional pre-mRNA splicing that we describe herein likely reflects the function of the canonical prefoldin complex.

It is known that coupling to transcription is not essential for splicing to occur (47). This can explain why we did not detect a possible contribution of prefoldin to pre-mRNA splicing under serum starvation conditions. We found that, in control cells, serum stimulation provoked increased efficiency of splicing, reflected in higher exon/intron ratios in total RNA reads. The transcriptional spur caused by serum stimulation might explain this phenomenon through co-transcriptional splicing. If that were the case, a positive contribution of prefoldin to splicing would be only expected under serum conditions, as we found.

Human prefoldin is involved in some cellular regulations like the epithelial-mesenchymal transition (EMT). PFDN1 has been described to mediate EMT in lung cancer cell lines and metastasis in mouse models (10). We have recently confirmed that overexpression of prefoldin subunits associates with the risk of mortality and metastasis in non-small cell lung cancer (48). The current model suggests that prefoldin overexpression would alter the expression pattern of EMT regulatory genes (10). The results of this study may contribute to elucidate how prefoldin can play such a regulatory function.

## DATA AVAILABILITY

The RNA-seq and ChIP-seq data in this article is available at GEO (https://www.ncbi.nlm.nih.gov/geo/) with accession number GSE155231 and GSE171466, respectively.

The ChIP-seq data of RNA pol II and Ser2P RNA pol II used in Figure 5 was obtained from GEO with accession numbers GSM1545657 and GSM4113565, respectively.

## ACKNOWLEDGEMENTS

We thank all the people of the IBiS gene expression lab for helpful discussion, Victoria Begley for English editing, and Cindy Will and Reinhard Lührmann for anti PRP19C antibodies. We thank all technicians from IBiS facilities, and E. Andújar and M. Pérez from the CABIMER Genomic Unit for technical assistance.

---

**Supplementary figure S1.**
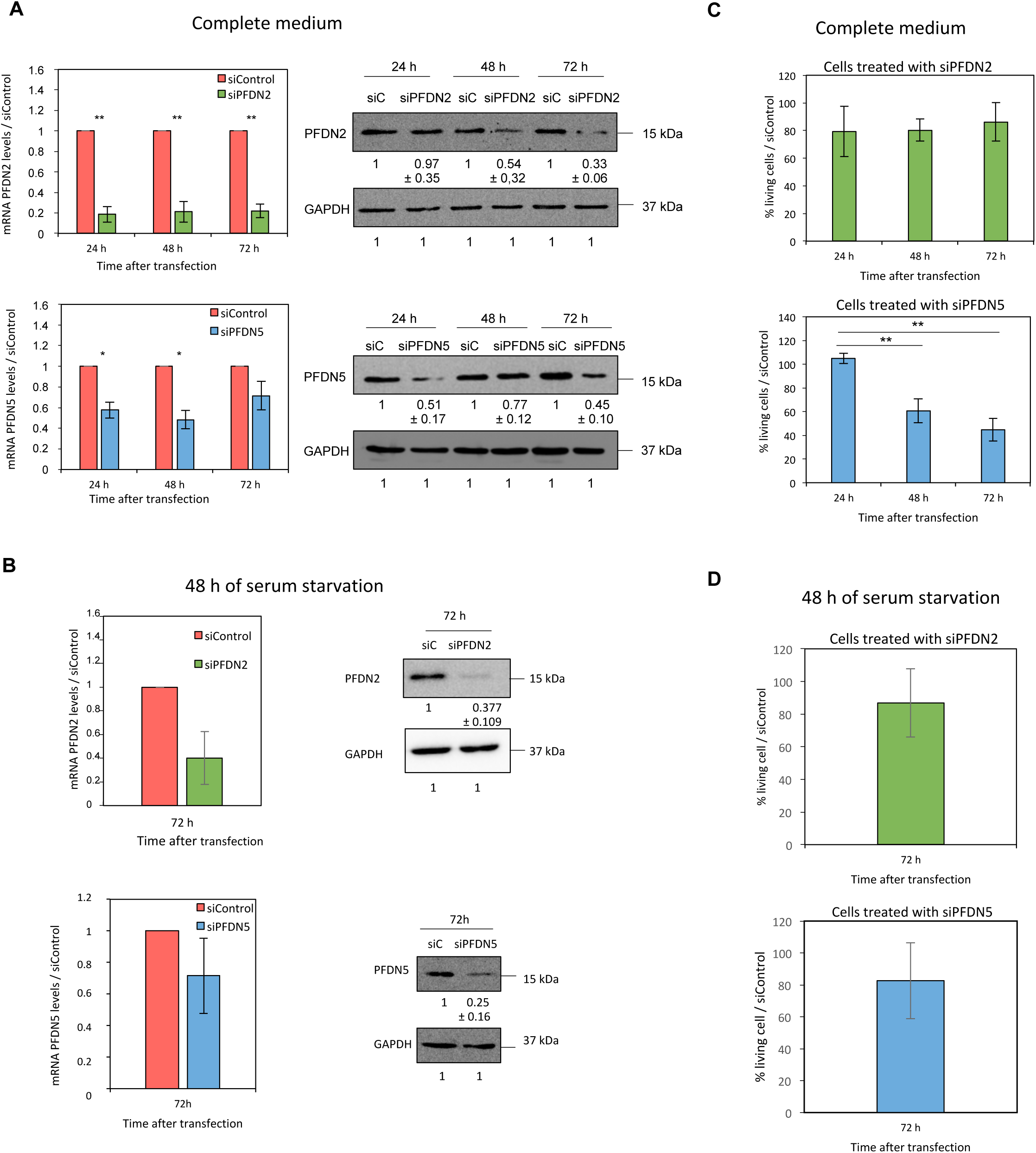
Efficiency of the siRNA-mediated depletion of PFDN2 and PFDN5. A) Left panels: PFDN2 and PFDN5 mRNA levels in HCT116 cells, 24, 48 and 72 hours after being transfected with siPFDN2 or siPFDN5, respectively, or with a siRNA control (siControl). Right panels: Protein levels analyzed by Western blotting using antibodies against PFDN2 or PFDN5 and GAPDH as a loading control, after transfecting the cells with siPFDN2, siPFDN5, respectively, or with siControl. The results were quantified with Image Lab software. B) Left panels: PFDN2 and PFDN5 mRNA levels in HCT116 cells treated with siPFDN2, siPFDN5, or siControl for 24 h and serum starved for 48 h more (72 hours total siRNA treatment). Right panels: Protein levels of the same samples analyzed by western blotting using antibodies against PFDN2 or PFDN5 and GAPDH as a loading control. C) Cell viability assay on the same cell line transfected with siPFDN2 or siPFDN5 and the siControl separately. The cells were counted every 24 hours for 3 days, and the percentage of cells relative to the control was represented. D) Cell viability assay in the same conditions as in B. In all panels, averaged values ± the standard deviation obtained from three different experiments are represented. * Student’s t test with p value < 0.05 and ** Student’s t test with p value < 0.005.

**Supplementary figure S2.**
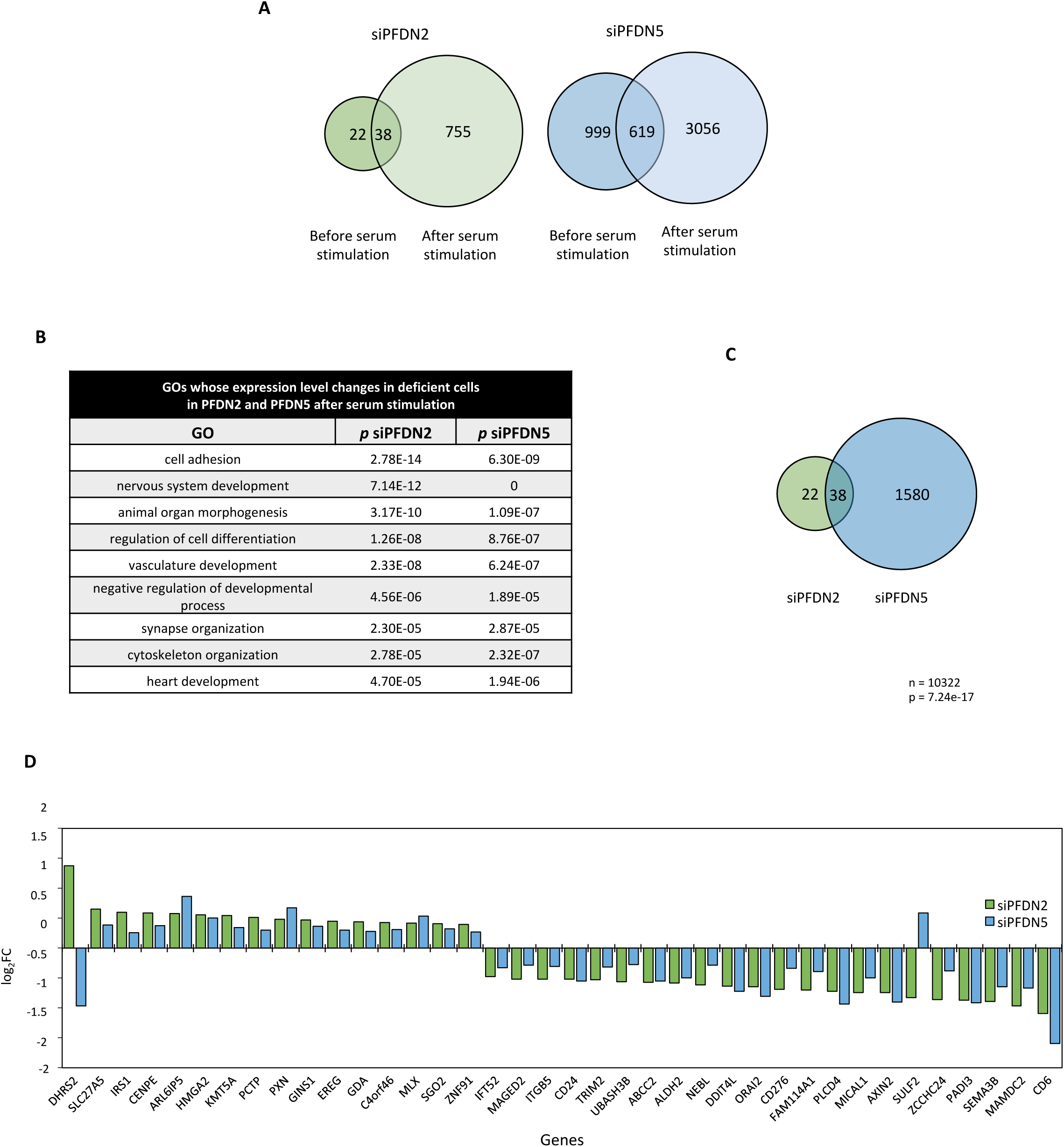
Differential expression of PFDN2- and PFDN5-deficient cells. A) Representation of the number of genes differentially expressed in the cells transfected withsiPFDN2 or with siPFDN5 at each time, before and after serum stimulation. B) The genes affected by siPFDN2 or siPFDN5 after serum stimulation were functionally grouped into Gene Ontology categories. Those categories that were commonly affected by both siRNAs are represented in the table. C) Representation of the number of genes differentially expressed in the cells transfected with siPFDN2 or with siPFDN5 before serum stimulation, and the genes that coincide. p value of the hypergeometric test is shown. n indicates total number of genes included in the analysis. D) Parallelism of the expression changes produced by siPFDN2 and siPFDN5. Changes in mRNA levels of genes affected by both siPFDN2 and siPFDN5 before serum stimulation are shown.

**Supplementary figure S3.**
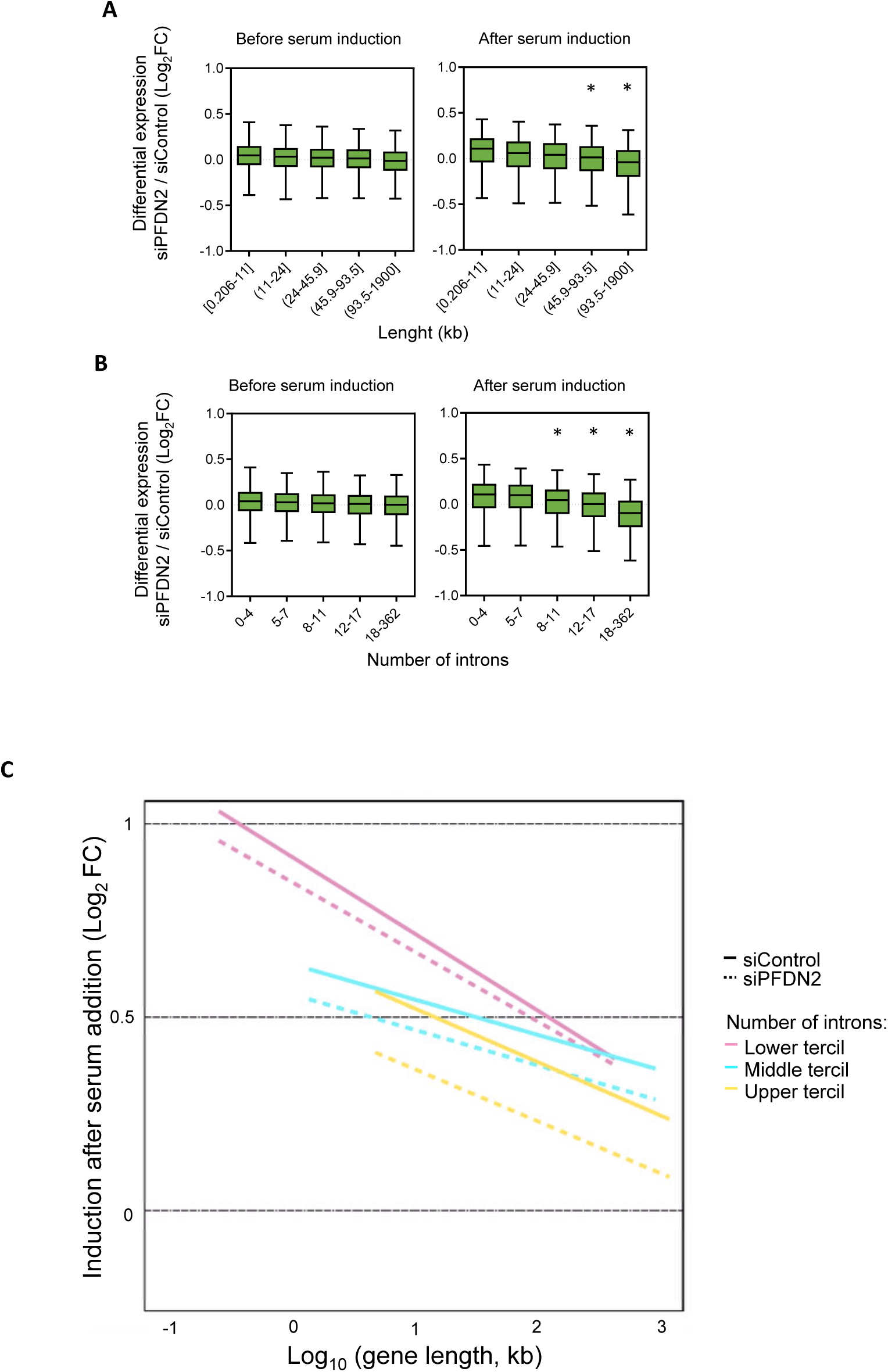
PFDN2 deficiency impairs the induction of long genes with a high number of introns. A) Expression differences between siPFDN2 and siControl cells (log2(FC)) with respect to the gene length, before and after serum stimulation. The genes were separated into quintiles. The * represents p < 0.005 in a Student’s t test comparing each quintile to quintile 1. B) The same analysis of A with respect to the number of introns that each gene contains, before and after serum stimulation. The genes were separated into quintiles. The * represents p< 0.005 in a Student’s t test comparing each quintile to quintile 1. C) Serum regulated genes (|log2(FC)| > 0 and FDR < 0.05, N = 1035) were divided into three groups according to the tercile of the number of introns. Serum-dependent fold change (Log_2_) is represented against the gene length (Log_10_), in cells transfected with the siControl (solid line) or siPFDN2 (dotted line).

**Supplementary figure S4.**
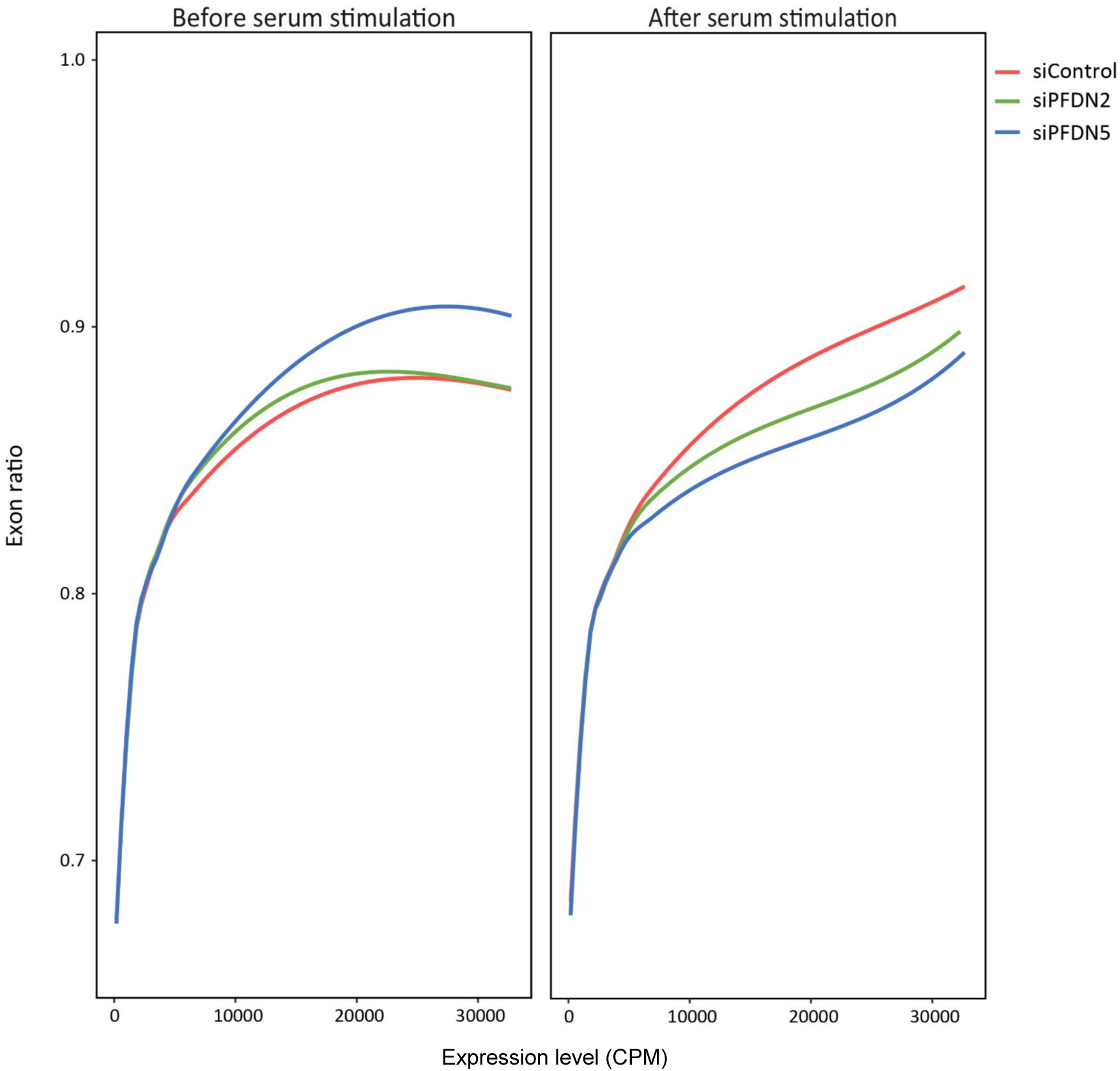
Exonic ratio distribution before and after serum stimulation. The exonic ratio (vertical axis) is represented against the level of gene expression (horizontal axis) of the cells transfected with the siControl (red), siPFDN2 (green) and siFDN5 (blue) before and after serum stimulation.

**Supplementary figure S5.**
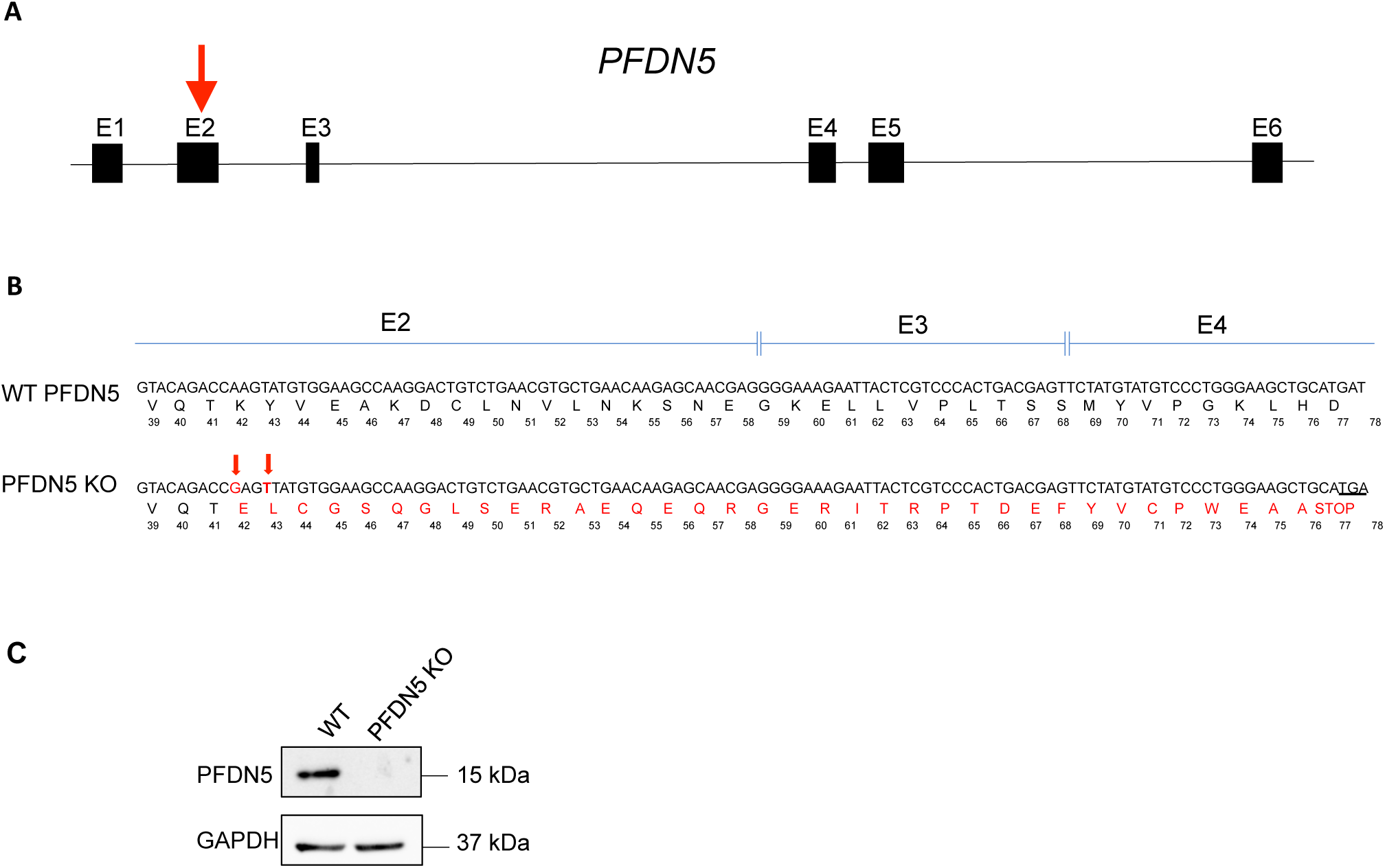
The sequence of the *PFDN5* KO. A) A frameshift mutation of *PFDN5* was originated by CRISPR-Cas9 technology in its exon 2. B) The sequence of the three exons affected by the translational consequences of the frameshift mutation is shown, comparing the WT and the *PFDN5* KO cell line. The bases in red represent the two mutations caused by CRISPR-Cas9 (an A to G transition and a T insertion, marked with arrows). Predicted translation is shown under each DNA sequence. The numbers under the aminoacid residues correspond to the position in the PFDN5 WT protein from its N-terminus. The residues in red are those not found in the WT protein in each position. A premature stop codon in exon 4 is underlined. C) Western blot showing the absence of PFDN5 protein in the *PFDN5* KO cell line.

**Supplementary figure S6.**
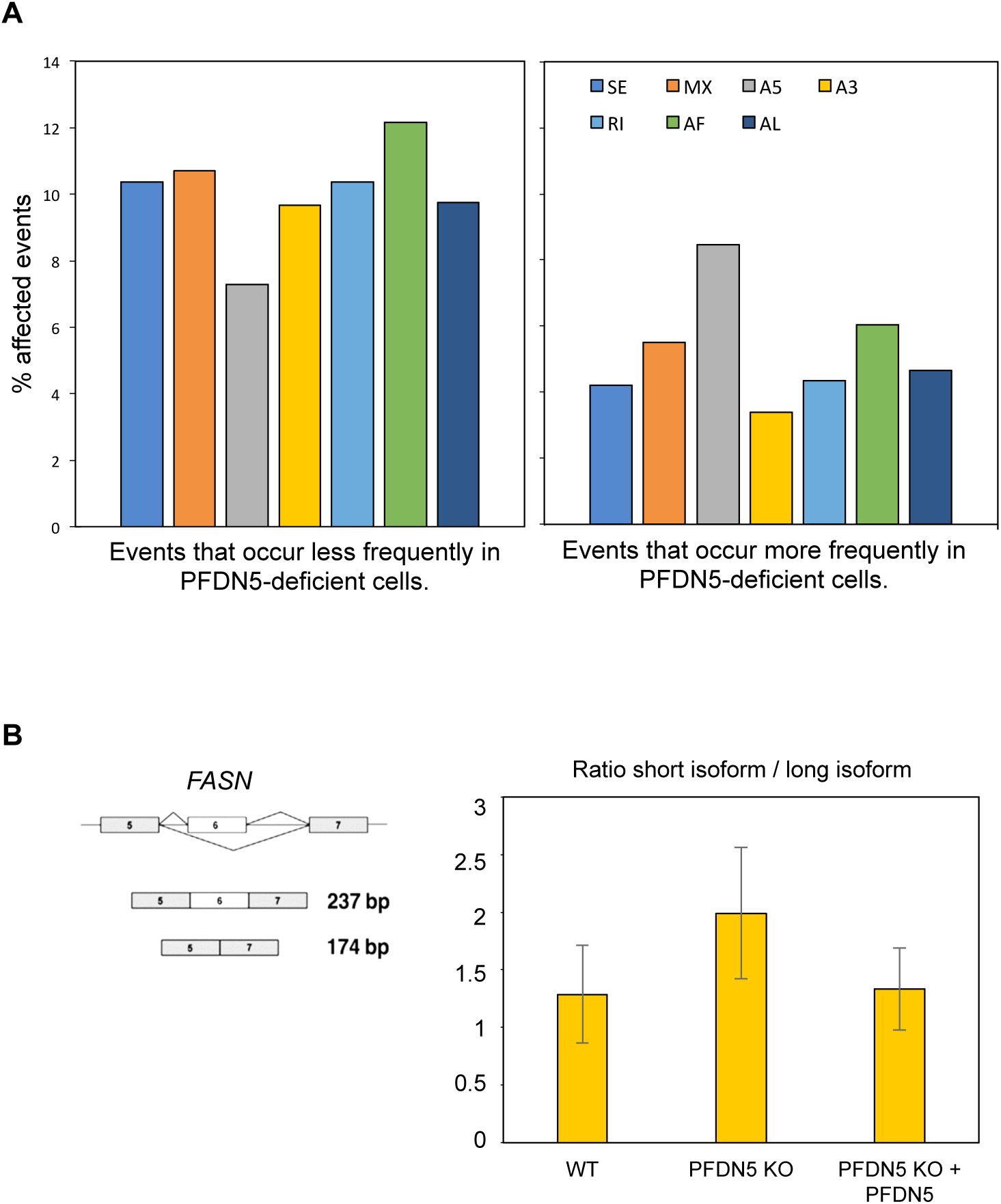
PFDN5 deficiency has a mild general impact on alternative splicing. A) PFDN5 depletion causes general alteration in alternative splicing. SUPPA computational tool (19) was used for exploring the impact of siPFDN5 on alternative splicing. Bars histogram representing the percentage of annotated alternative splicing events that occur significantly less frequently in PFDN5-deficient cells is shown in the left panel. On the right, bars histogram representing the frequency of alternative splicing events that occur more frequently in PFDN5-deficient cells. SE: skipping exon; MX: mutually exclusive exons; A5: alternative 5’ splice-site; A3: alternative 3’ splice-site; RI: retained intron; AF: alternative first exon; AL: alternative last exon. B) Lack of PFDN5 causes accumulation of intron reads after serum stimulation in a *FASN* minigene assay. A schematic drawing of the alternative isoforms of the Fas minigen is shown on the left. On the right, bar histogram of short isoform / long isoform ratios of the Fas minigen, in control cells, *PFDN5* KO cells, and *PFDN5* KO cells transfected with GFP-PFDN5. The graph represents the average values and the standard deviation obtained from five different experiments. According to the Student’s t test, the difference between WT and KO cells and between KO and KO+GFP-PDFN5 cells did not generated p values lower than 0.05.

**Supplementary figure S7.**
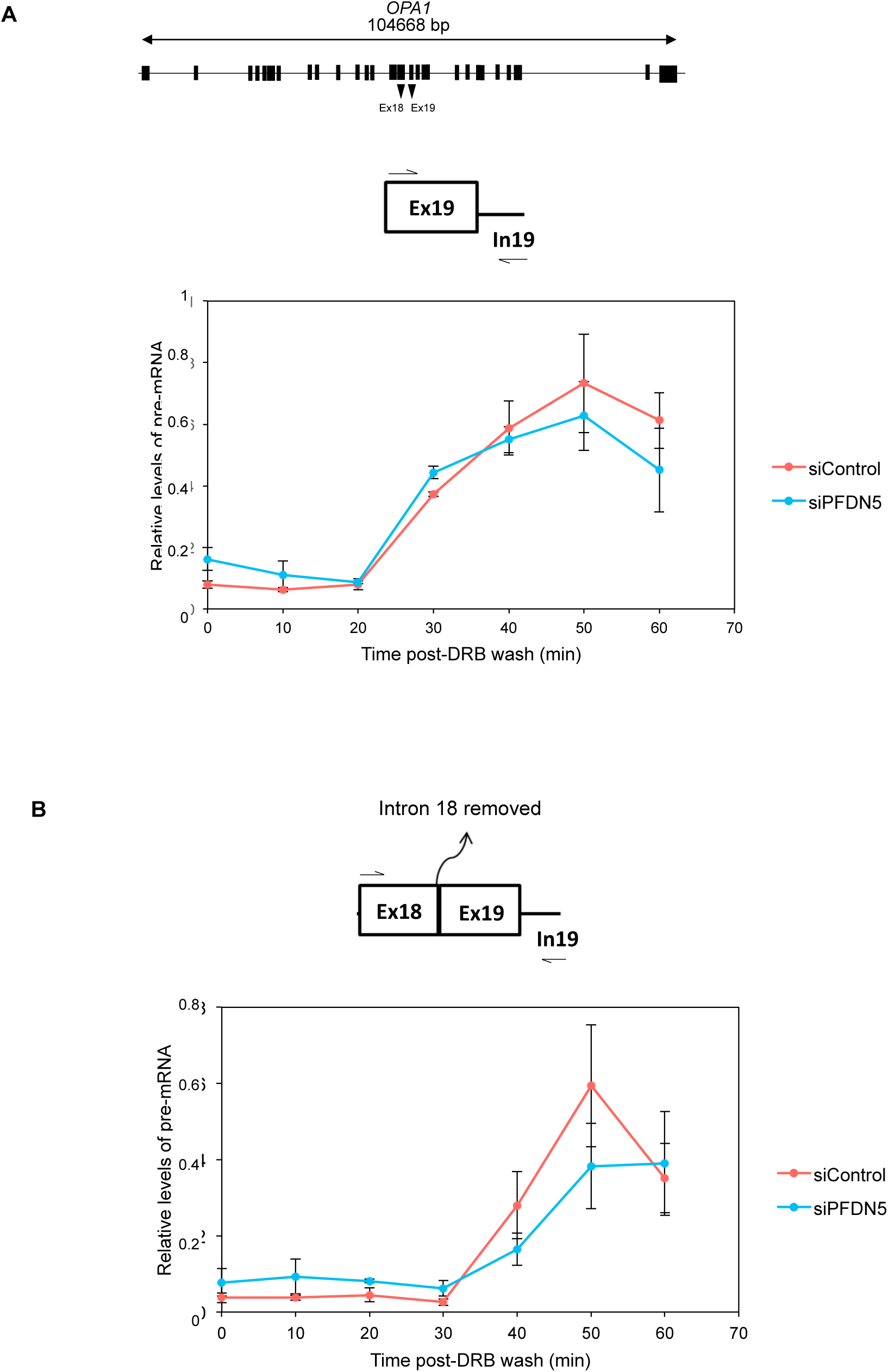
PFDN5 depletion impairs co-transcriptional splicing in the long *OPA1* gene. **A)** Pre-mRNA levels from the *OPA1* gene in cells transfected with control siRNA (red line) or PFDN5 siRNA (blue line). Cells were treated with 100 µM DRB for 3 hours. Samples were taken every 10 minutes after washing DRB out. Pre-mRNA data was normalized first to the mature mRNA, and next to the DRB untreated sample. A scheme of the *OPA1* gene is also shown, and the amplicon used is depicted above the graph. **B)** Event of co-transcriptional splicing (measured as the presence of pre-mRNA without the indicated intron) in the *OPA1* gene in cells treated as in A). The amplicon used is depicted above the graph. The average and standard error of at least three biological replicates with three technical replicates each are represented.

**Supplementary figure S8.**
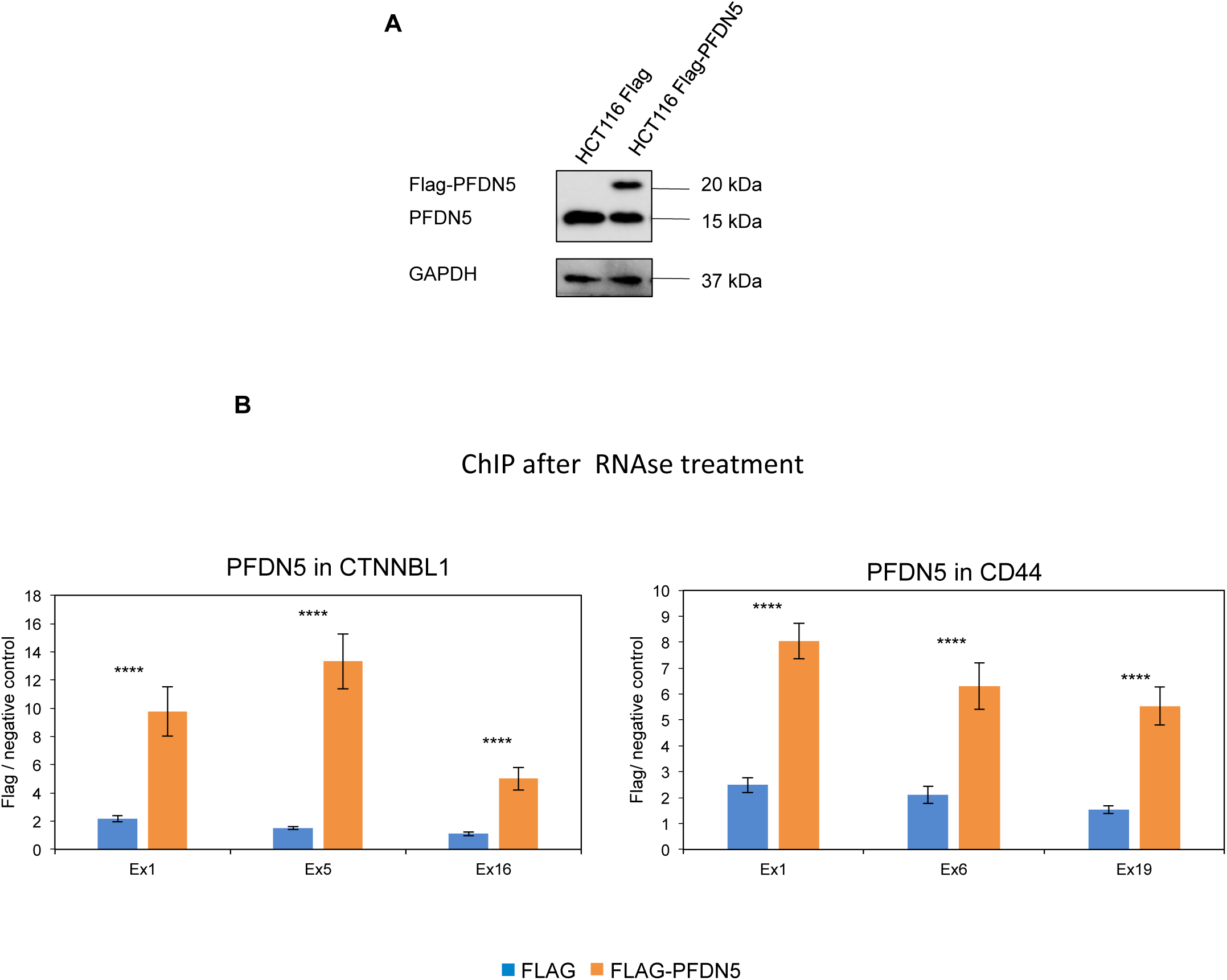
PFDN5 presence in the chromatin is not RNA dependent. A) HCT116 cell line expressing Flag-PFDN5. Expression of Flag-PFDN5 in HCT116 stable cell line. A western blot is shown, after using antibodies against PFDN5 and GAPDH as a loading control. B) The level of Flag-PFDN5 in the chromatin at different loci throughout the *CTNNBL1* gene (left panel) and the *CD44* gene (right panel), measured after treatment of the extracts with RNase. The graphs represent the average values ± the standard error obtained from three different experiments. ****Student’s t test with p value < 0.0001. A total number of 12 technical replicates were considered for Student’s t test.

**Supplementary figure S9.**
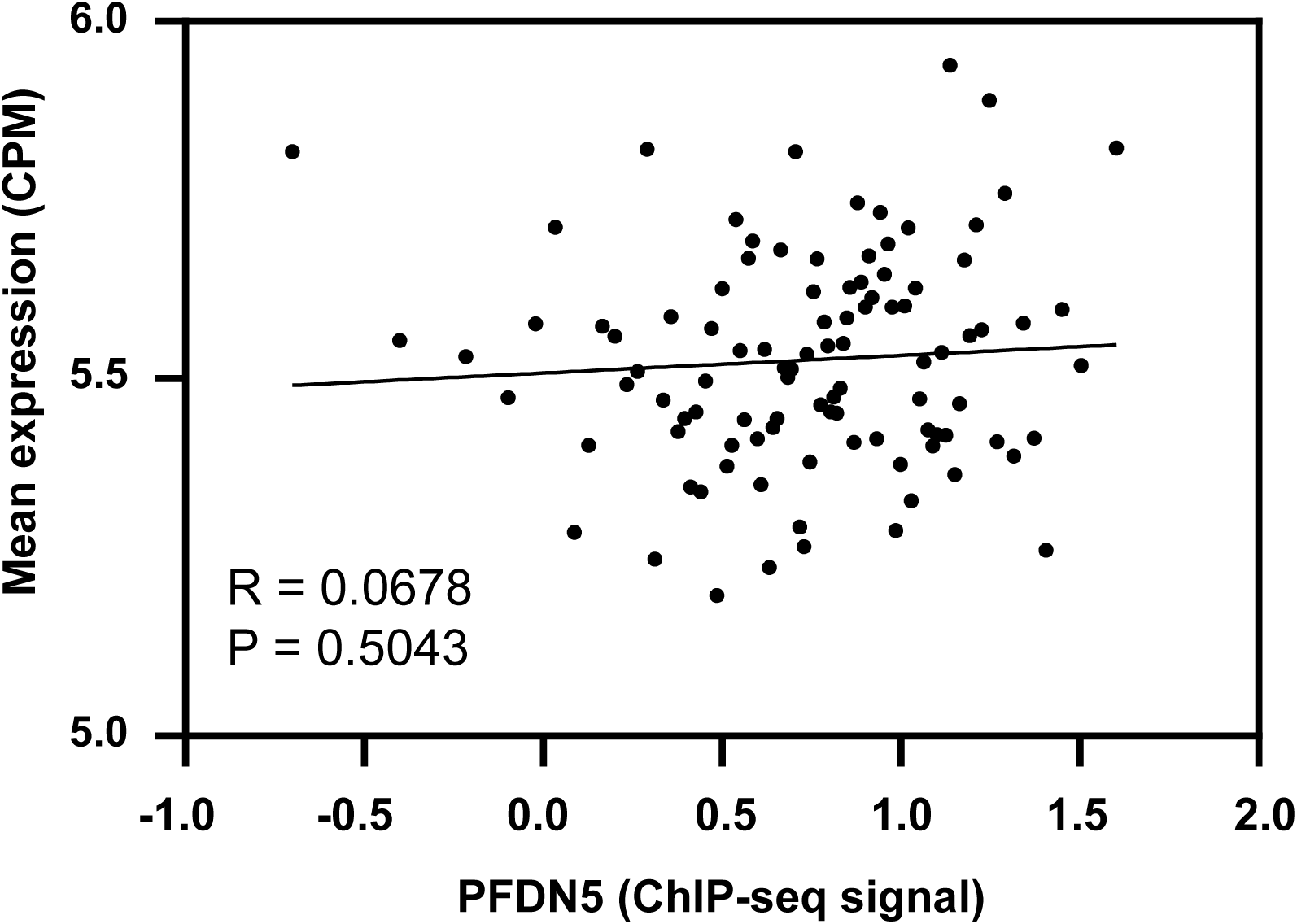
Weak correlation between prefoldin ChIP-seq signal and total gene expression. The promoter (TSS +/- 500 bp) signal of PFDN5 was represented with respect to the level of gene expression. Expressed genes (>0.1 reads per kilobase per million, RPKM) were ordered according to the level of expression and divided into 100 bins. The mean signal of each group was the calculated and represented. The R and p values from Pearsońs correlation are shown.

**Supplementary figure S10.**
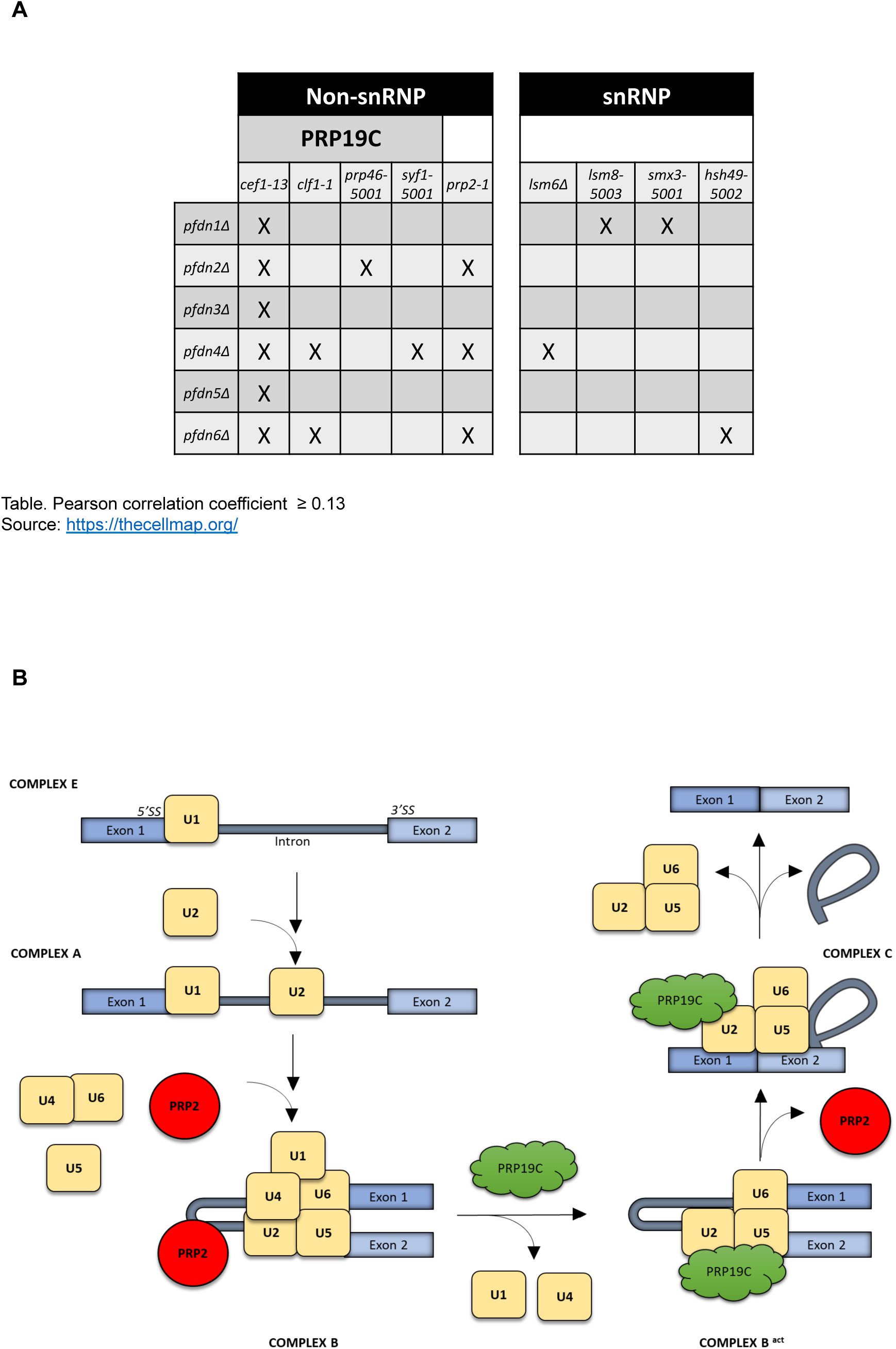
Correlations between prefoldin and splicing mutants in the CellMap database of genetic interactions. A) Genetic correlations between prefoldin and splicing mutants were explored in the Cell Map database (thecellmap.org). Pairs of prefoldin/pre-mRNA splicing mutants showing Pearson correlations coefficients higher than 0.13 are shown in the table. B) The two factors that showed general correlations with prefoldin mutants, PRP19C and PRP2, participate in the activation step of pre-mRNA splicing. Spliceosome assembly and splicing reaction. snRNPs subunits U1, U2, U4, U5 and U6 are represented; as well as the splicing factors Prp2 and Prp19C.

**Supplementary figure S11.**
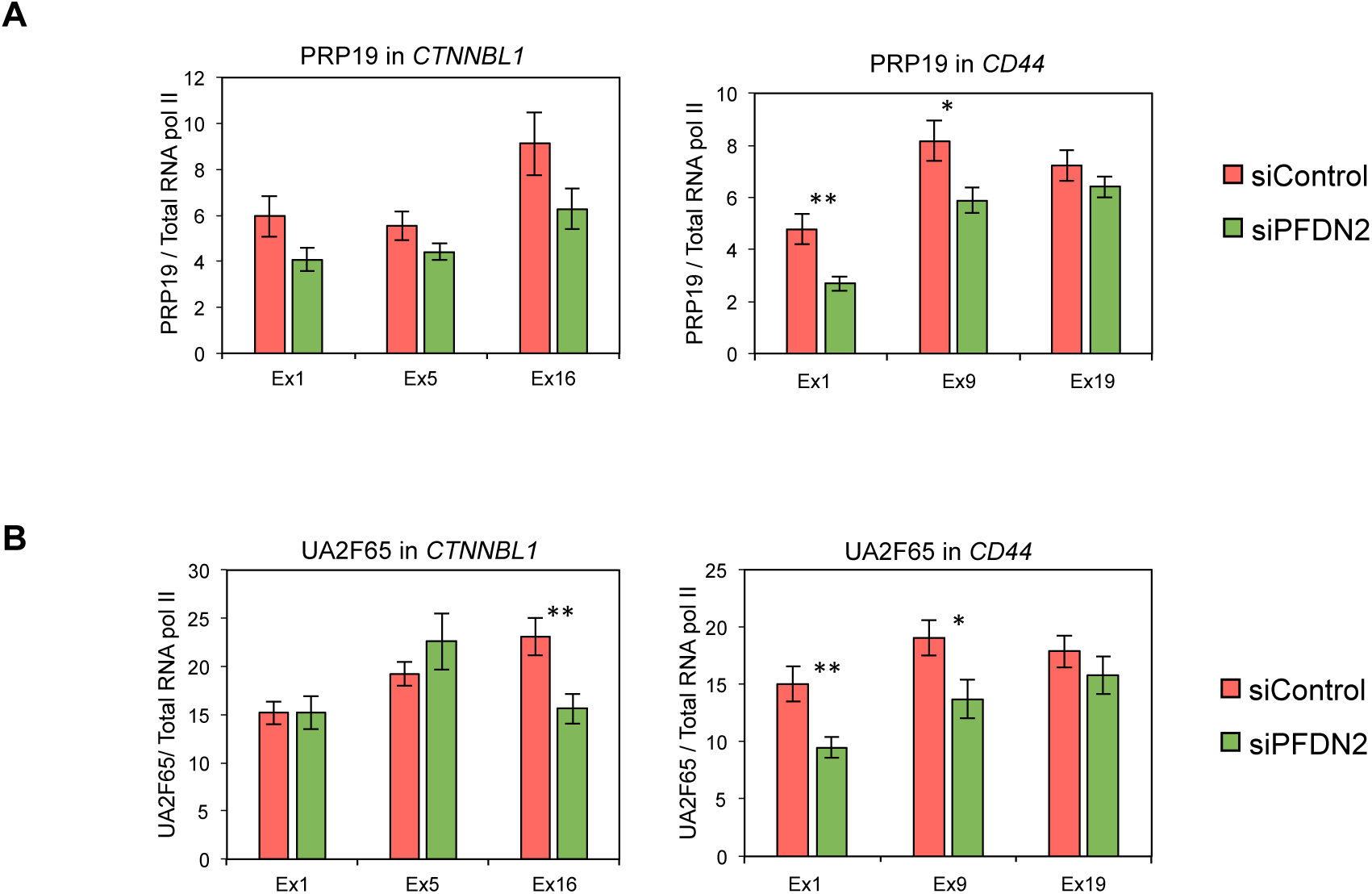
Splicing factors recruitment is impaired in PFDN2 knockdown cells. A-B) The levels of PRP19 (A) and U2AF65 (B) were measured by ChIP in different regions of the *CTNNBL1* gene (left panels) and the *CD44* gene (right panels) in cells treated with siControl (red bars) and with siPFDN2 (green bars). In all the panels, values were normalized to total RNA pol II levels. The graphs represent the average values ± the standard deviation obtained from three different experiments. * p value < 0.05; ** < 0.005. A total number of 12 technical replicates were considered for Student’s t test.

**Supplementary figure S12.**
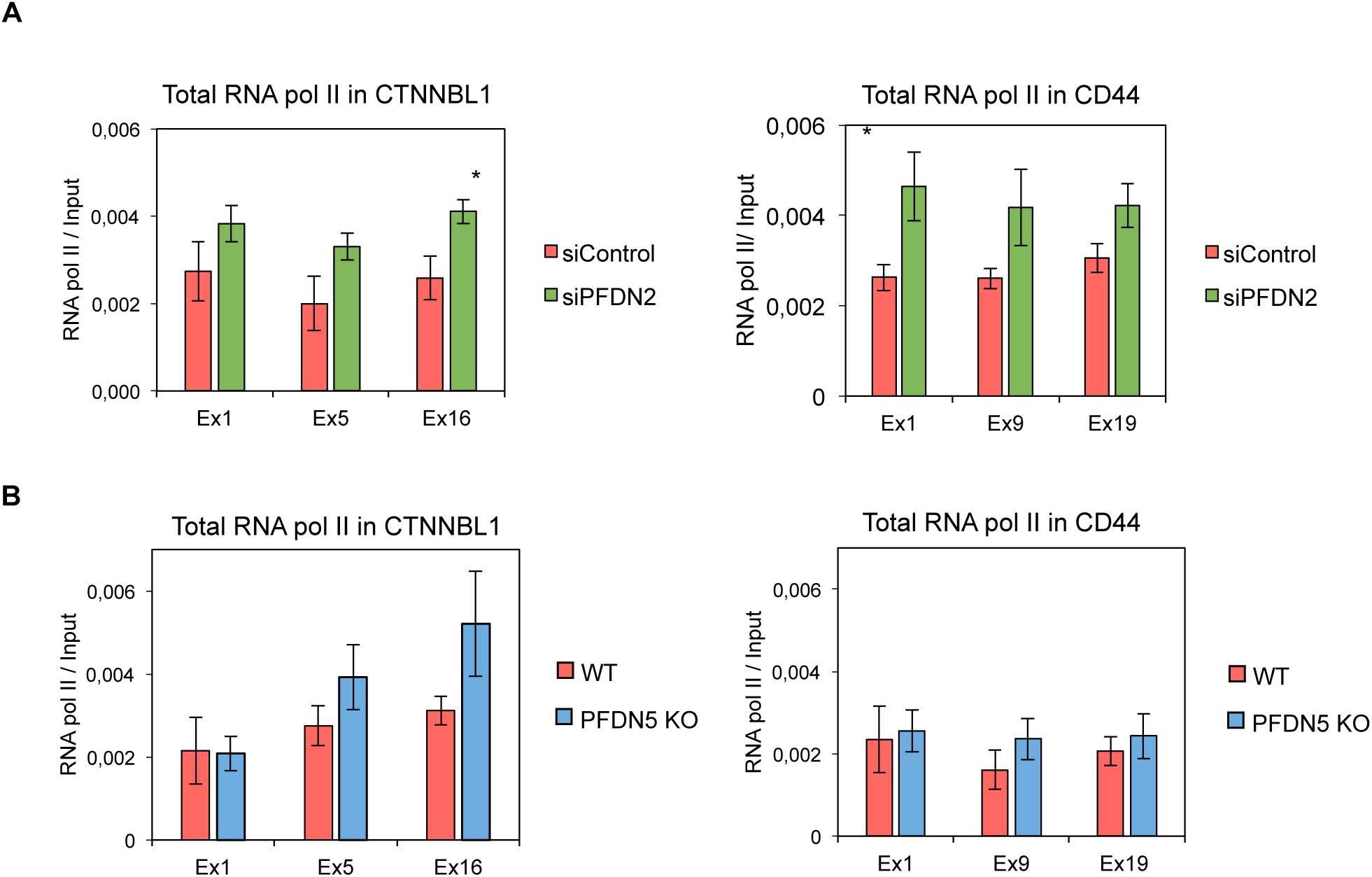
Total levels of RNA pol II remain constant in the absence of prefoldin. A and B) The levels of RNA pol II were measured by ChIP in different regions of the *CTNNBL1* gene (left panels) and the *CD44* gene (right panels), using the 8WG16 antibody in A) cells treated with siControl (red bars) and with siPFDN2 (green bars); B) control (red bars) and *PFDN5* KO cells (blue bars). The graphs represent the average values ± the standard deviation obtained from three different experiments. * p value < 0.05. A total number of 12 technical replicates were considered for Student’s t test.

**Supplementary figure S13.**
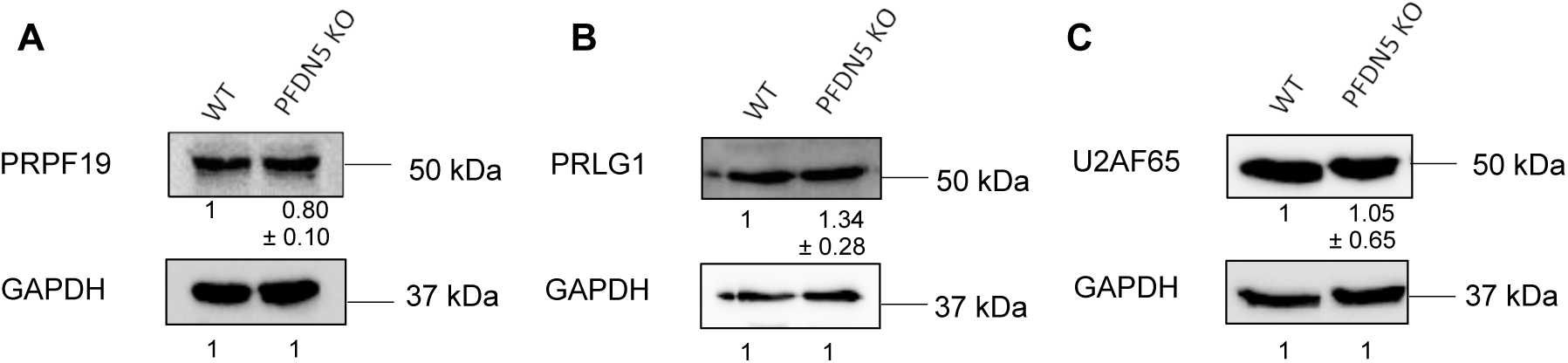
Total levels of different splicing factors and RNA pol II remain constant in the absence of prefoldin. A-C) Protein levels determined by western blotting, using antibodies against PRPF19 (A), PRGL1 (B) and U2AF65 (C) splicing factors and GAPDH as a loading control. The results were quantified with Image Lab software. Averaged values ± the standard deviation obtained from three different experiments are represented.

**Supplementary figure S14.**
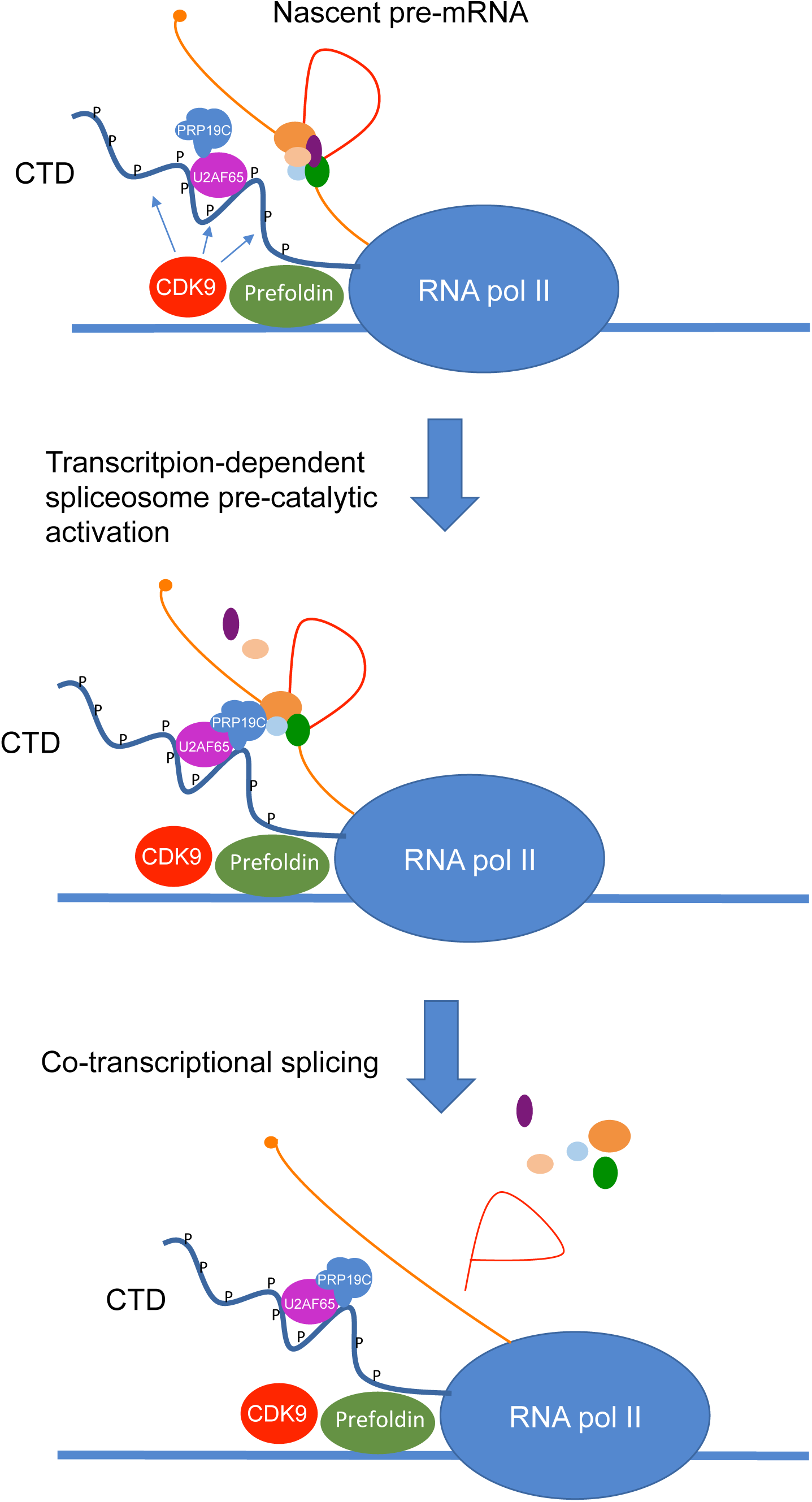
Graphic model illustrating how prefoldin could contribute to co-transcriptional splicing. Prefoldin helps recruit CDK9 to transcribed chromatin. CDK9 phosphorylates the RNA pol II CTD at Ser2 residues. In turn, the phosphorylated CTD recruits U2AF65 and the PRP19 complex. These splicing factors, acting from the CTD, activate the spliceosome assembled on the nascent pre-mRNA, thereby favouring co-transcriptional pre-mRNA splicing.

**Supplementary figure S15.**
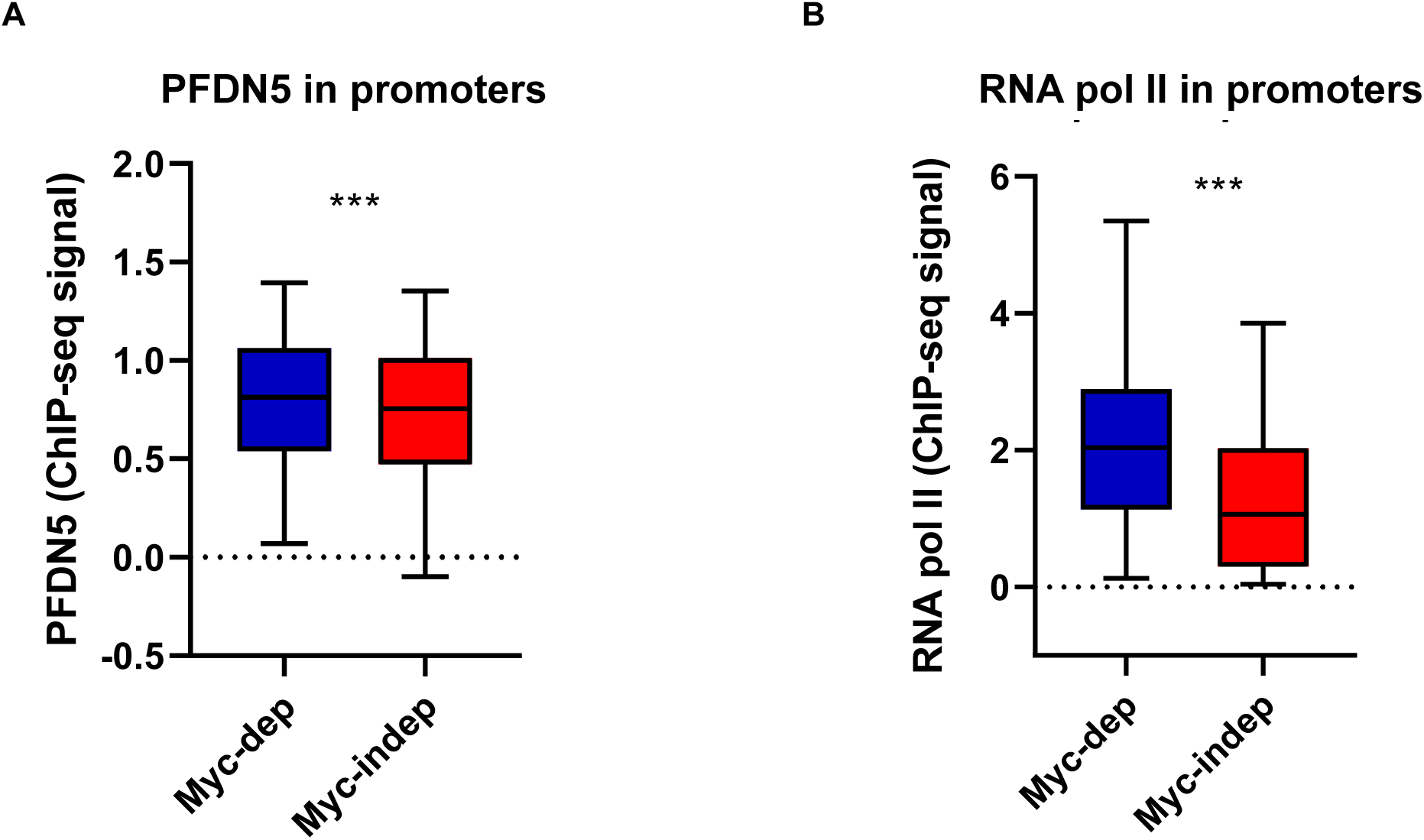
PFDN5 binds both Myc-dependent and -independent genes. A) Box plot comparing the presence of PFDN5 in promoters of Myc-dependent or independent genes. Data from ChIP-seq analysis were used. B) Box plot comparing the presence of RNA pol II in promoters of Myc-dependent or independent genes. *** Mann-Whitney U test p value < 0.0005.

**Supplementary Table S1.**
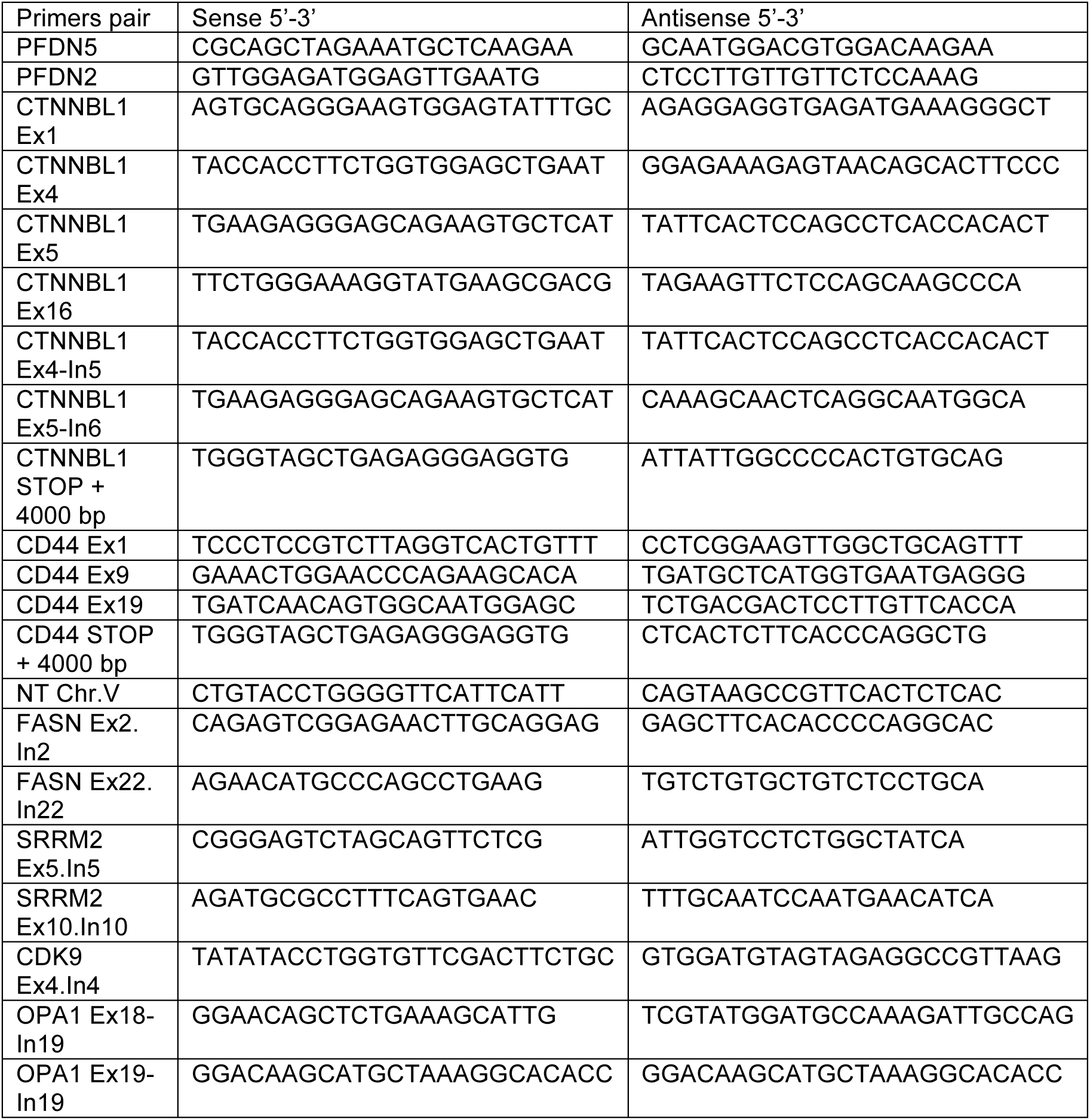
Primers used in this work.

